# High-throughput identification of chromatin-opening transcription factors

**DOI:** 10.64898/2026.10.07.757445

**Authors:** Aidan Keith, Shruti Jain, Riza M. Daza, Chengxiang Qiu, Pierre-Luc Germain, Jay Shendure, Silvia Domcke

## Abstract

Cell fate transitions involve major changes to the chromatin landscape, some of which are thought to deepen the basin around a newly adopted cell state, stabilizing gene expression programs that might otherwise be transient. Transcription factors (TFs) are central to this process, binding only a subset of the genome-wide occurrences of their cognate motifs to create accessible regions that can act as cell type-specific enhancers. However, apart from a few well-studied cases, it is not clear which of the ∼1,600 mammalian TFs can directly or indirectly initiate newly accessible sites, nor why only some motif occurrences gain accessibility. Here, we performed pooled CRISPR activation (CRISPRa) screens to upregulate endogenous developmental TFs in mouse embryonic stem cells (mESCs) and assess their impact on the chromatin landscape. Following a pre-screen of 240 developmental TFs for their capacity to elicit strong changes to the transcriptome, we prioritized 60 TFs for a well-powered CRISPRa multiome screen in which we jointly profiled gene expression and chromatin accessibility across ∼140,000 single cells. We identify TFs that appear to initiate new chromatin accessibility on their own (“autonomous”), others whose chromatin-opening activity appears dependent on coincident binding of co-TFs heterogeneously expressed across mESCs (“opportunistic”), and yet others which appear to induce a co-TF and then act alongside it (“hierarchical”). Taken together, our findings suggest that the ability of a TF to “pioneer” chromatin opening, when it is newly expressed at physiological levels, strongly depends on the cell state and gene regulatory network it lands in. Looking forward, a comprehensive charting of such dependencies will inform how cell fate transitions are initiated in development, homeostasis and disease, and how to engineer them *in silico, in vitro* and *in vivo*.

## INTRODUCTION

Transcription factors (TFs) bind *cis*-regulatory elements (CREs) in accessible regions of the chromatinized genome to govern target gene expression. During cell state transitions, new CREs become accessible, but the rules governing which sites open, and in which cell states, remain largely unknown. Proposed mechanisms by which TFs access closed regions include cooperativity between TFs and chromatin remodelers, opportunistic binding during nucleosomal turnover, and direct displacement of nucleosomes by “pioneer factors”^1–3^. However, for nearly all mammalian TFs, three basic questions remain open: (i) whether they can initiate accessibility changes at all, directly or indirectly; and, for those that can, (ii) whether that capability is contingent on cellular context; and (iii) what determines which motif occurrences become accessible.

In principle, these questions can be answered by initiating expression of a TF in defined cell states and measuring genome-wide changes in accessibility, thus quantifying the TF’s capacity to modulate chromatin and accruing the positive and negative examples needed to model binding site selection. To date, however, such experiments have largely been performed one TF at a time, *e.g.* overexpressing the TF and profiling by bulk ATAC-seq and/or ChIP-seq^2,4,5^. Pooled single-cell screens, in which each cell newly expresses a distinct TF, overcome this bottleneck^6,7^, but when read out by single-cell RNA-seq (scRNA-seq), forgo the accessibility measurement that would inform each TF’s capacity for chromatin-opening. Single-cell co-assays of accessibility and gene expression (sc-multiome) are a promising solution, but are costly to scale. To date, with few exceptions^8^ most sc-multiome TF perturbation screens either do not call differentially accessible sites^9^, or assess only TF knockouts and knockdowns and thus speak to accessibility maintenance but not the initiation of new sites^10–13^.

In any “TF activation” screen, one must decide both how and how strongly to express each TF. Most prior studies use open reading frame (ORF) overexpression^6,9^, well-suited for driving cell state reprogramming. But ORF overexpression can reach supraphysiological levels at which motif usage becomes progressively indiscriminate^14–16^, and often lack 3’ UTRs shown to be essential for protein activity^17^. An alternative is CRISPR activation (CRISPRa), in which a catalytically dead Cas9 fused to a transcriptional activator drives expression from an endogenous promoter^18,19^. Although CRISPRa yields lower and less consistent levels of activation, it operates within a more physiologically reasonable range, and may be more likely than pre-synthesized ORFs to recapitulate endogenous isoform choice -- a potentially critical difference, as two-thirds of alternative TF isoforms differ substantially in activity from the reference isoform^20^.

Here we sought to ask which developmentally relevant TFs, when newly expressed in naive pluripotent stem cells at physiologically reasonable levels, can open chromatin and alter gene expression programs, as well as whether such capabilities exhibit *cis*- or *trans*-dependencies. To this end, we first confirmed that CRISPRa of individual TFs yields motif-enriched accessibility gains in mESCs. Next, we evaluated which of 240 developmentally relevant TFs substantially altered the transcriptome when targeted by CRISPRa. Finally, we prioritized 60 TFs that drove the strongest transcriptional changes and performed a well-powered CRISPRa sc-multiome screen on these, profiling both gene expression and chromatin accessibility across ∼140,000 single cells. A subset of these TFs stood out as driving striking gains in accessibility. Furthermore, we could distinguish between: (i) TFs that appear to act on their own (“autonomous”); (ii) TFs whose chromatin-opening activity appears dependent on coincident binding of co-TFs heterogeneously expressed across mESCs (“opportunistic”); and (iii) TFs that appear to induce a co-TF and then act alongside it (“hierarchical”). Beyond nominating new TFs with chromatin-opening potential, our results suggest that the capacity of a TF to open chromatin and alter gene expression is highly context-dependent, and demonstrate a cost-effective path towards systematically charting such dependencies.

## RESULTS

### CRISPRa of unexpressed TFs alters chromatin accessibility in mESCs

We generated a monoclonal mESC line stably expressing dCas9-VPR^21^ and used qRT-PCR over a five-day time course to identify a suitable timepoint for our screens (**Supplementary Fig. 1A**). Because we targeted TFs that are not detectably expressed in mESCs, activation was quantified relative to a highly expressed reference (*Gapdh*) rather than to baseline TF expression. CRISPRa of three TFs (*Fosl1*, *Gata6*, *Neurog2*) yielded activation 10- to 30-fold lower than transduction of cDNA constructs under a strong promoter, but on par with *Gapdh*, confirming their successful activation (**Supplementary Fig. 1B-C**). As activation was consistently high 96 hrs after transduction and completion of drug selection, we selected this timepoint for all downstream experiments.

To evaluate whether CRISPRa-activated TFs alter chromatin accessibility, we generated bulk ATAC-seq profiles of gRNA- and cDNA-transduced populations. Both modes of activation revealed differentially gained (DG) and differentially lost (DL) sites (**Supplementary Fig. 1D**). DG sites substantially overlapped between the two methods of TF activation, and the top enriched motif among DG sites exhibiting the highest fold-changes (FC) consistently corresponded to the targeted TF (**Supplementary Fig. 1E-H**). There were more DL sites when TFs were overexpressed from cDNAs rather than activated by CRISPRa (**Supplementary Fig. 1D**). These DL sites were enriched for OCT4 and SOX2 motifs, suggesting that supraphysiological TF levels may drive exit from pluripotency (**Supplementary Fig. 1F-H**).

To ask whether such changes are detectable in pooled single-cell screens, we targeted 10 TFs with CRISPRa, including 7 previously reported to have pioneer activity (*Gata6, Neurog2, Ascl1, Otx2, Ebf1, Pax7, Onecut1*)^22–28^ and 3 reported to lack it (*Pou3f2/Brn2, Tbx19/Tpit, Pax5*)^24,29,30^, alongside non-targeting control (NTC) gRNAs. At 96 hrs post-transduction, we performed single-cell ATAC/RNA multiome profiling (10x Genomics). Following QC, we obtained profiles for 11,615 cells (median 3,184 RNA UMIs and 10,860 unique ATAC fragments per cell; **Supplementary Fig. 2A-B**). Of these, 6,433 (55%) had at least one assigned gRNA (**Supplementary Fig. 2C-D**). Counting cells with one or more assigned gRNAs, a median of 831 cells was recovered per target TF; 797 cells were assigned a single NTC gRNA. Target TF expression was elevated in cells bearing the corresponding gRNA, confirming on-target activation (**Supplementary Fig. 2E**).

TF-driven changes in chromatin accessibility were evident in these single-cell data as well. For example, the 304 cells bearing *Gata6* gRNAs, when pseudobulked, bore 3-fold more DG sites than the 797 NTC cells, and ∼40% of these sites overlapped those gained in a non-pooled experiment (*i.e.* CRISPRa of *Gata6* in mESCs → bulk ATAC-seq) (**Supplementary Fig. 3A-B**). GATA was the top enriched motif in the overlapping sites, while FOS/JUN was the top motif in the non-overlapping sites, possibly reflecting greater differentiation of mESCs in the single-cell experiment and highlighting the value of pooled screens in which controls reside in the same dish (**Supplementary Fig. 3C**). Aggregating across DG sites identified by bulk ATAC-seq of CRISPRa-*Gata6* mESCs, pseudobulked CRISPRa-*Gata6* single cells exhibited elevated accessibility, while *Neurog2* gRNA(+) cells from the same experiment did not (**Supplementary Fig. 3D**). Across the full panel, among cells with detected target TF expression, 5 of 7 TFs reported to have pioneer activity showed motif-concordant gains in accessibility (*Gata6, Neurog2, Ascl1, Otx2, Onecut1*), whereas 2 of 3 TFs reported to lack it did not (*Tbx19, Pax5*) (**Supplementary Fig. 3E**).

Together, these pilot experiments demonstrate that a pooled CRISPRa screen coupled to scATAC-seq can identify motif-concordant gains in chromatin accessibility. However, they also underscore that the sparsity of scATAC-seq data demands many cells per target. This motivated us to develop a hybrid computational-experimental strategy to prioritize a subset of developmentally relevant TFs on which to focus.

### A transcriptional pre-screen of 240 developmental TFs

To identify a candidate list of developmentally relevant TFs to target with CRISPRa, we leveraged a curated ontogeny of developmental cell states spanning mouse development from zygote to pup^31^, extracting 531 TFs that are differentially expressed between pairs of connected cell states. After excluding TFs that lacked a known motif in either the HOMER or JASPAR databases, as well as those already highly expressed in mESCs, we were left with 240 candidate developmental TFs (**Fig. 1A**).

**Figure 1.**
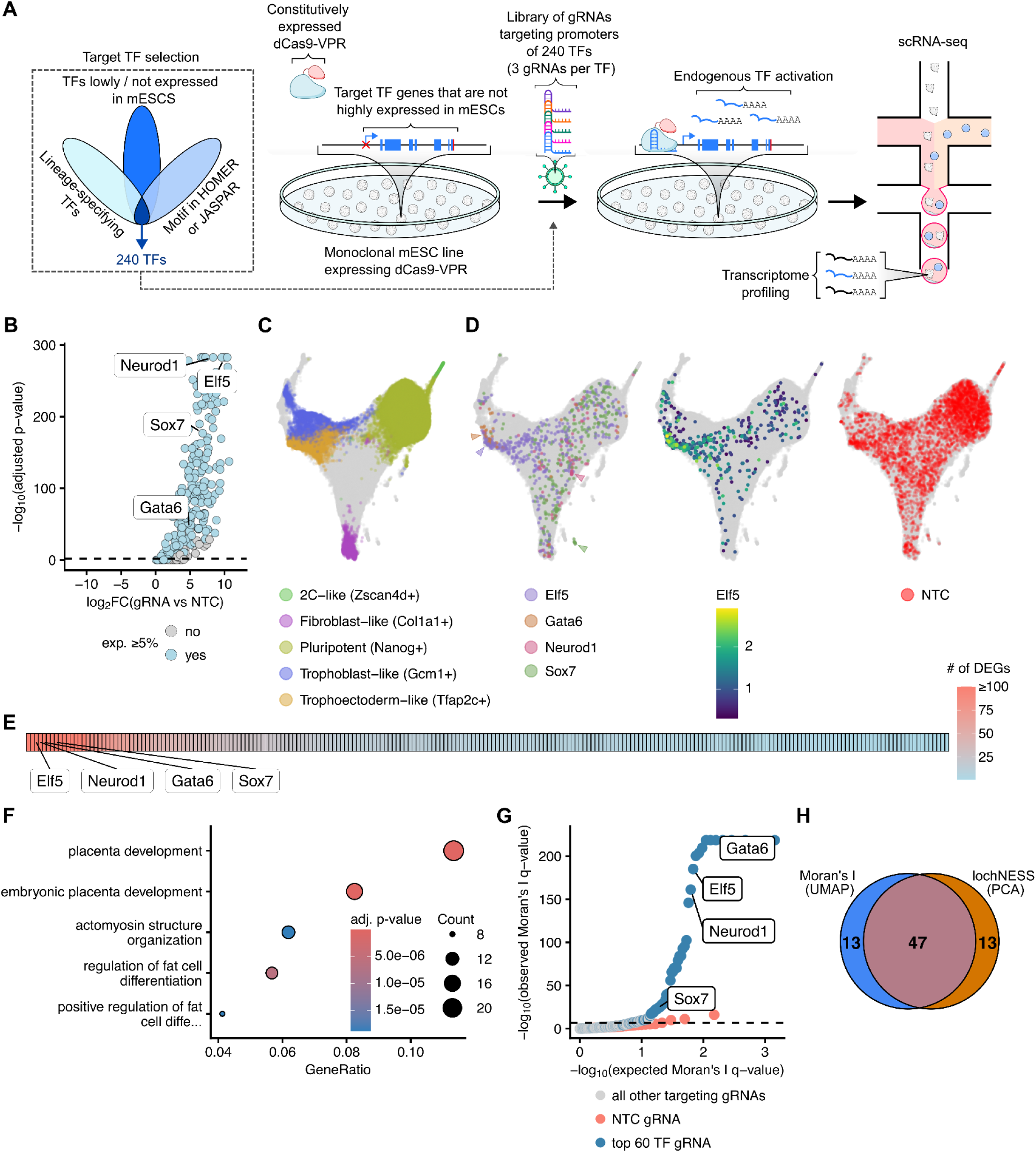
A transcriptional pre-screen of 240 developmental TFs. **(A)** Experimental schematic. **(B)** Volcano plot of differential gene expression of 240 TFs, comparing cells bearing TF-targeting vs NTC-targeting gRNAs (x-axis: log_2_ fold-difference, y-axis: -log_10_ Wilcoxon, adjusted p-values). Blue points indicate TFs with detected expression in ≥5% of cells bearing TF-targeting gRNAs. **(C)** UMAP embedding of scRNA-seq data, highlighting cells in which a given marker gene was both detected and locally enriched among its neighbors. Requiring both criteria suppresses sporadic expression, facilitating visualization of focal expression domains in a single embedding: *Nanog* (pluripotent cells), *Col1a1* (fibroblast), *Zscan4d* (2C-like), *Tfap2c* (trophectoderm-like) and *Gcm1* (trophoblast-like). For local enrichment, we required a cell’s lochNESS score^39^ to exceed the mean across cells in which the target TF was detected, or three times the mean for *Tfap2c* due to its broader expression pattern. **(D)** Same embedding as **C**, here highlighting cells with detection of gRNAs targeting *Elf5*, *Gata6*, *Neurod1* or *Sox7* (left), *Elf5* expression (middle), or NTC-targeting gRNAs (right). **(E)** Ranked heatmap of number of DEGs for each targeted TF (Wilcoxon adjusted p-value <0.01, expression detected in ≥5% of target cells, and fold-change relative to NTC ≥1.2). Selected TFs are labeled. **(F)** Dot-plot of GO Biological Process terms significantly enriched among the top 200 DEGs (ranked by FC) for *Elf5*; only the top 5 terms by adjusted p-value are shown. **(G)** Quantile-quantile (Q-Q) plot of expected vs observed Moran’s I q-values for targeting (blue) and NTC (red) gRNAs. Filled blue points highlight gRNAs targeting TFs chosen for the sc-multiome screen. **(H)** Venn diagram of the overlap (n = 47) between the top 60 TFs ranked by Moran’s I on the UMAP embedding in **C-D** and the top 60 ranked by gRNA-level lochNESS enrichment in PCA space (mean score across assigned cells, weighted by the number of assigned cells). Note that lochNESS is used in two distinct ways in this figure: to enhance visualization of marker-gene neighborhoods in **C**, and as a ranking statistic over gRNA assignments in **H**.

To prioritize further, we performed a primary screen of these 240 TFs with CRISPRa and scRNA-seq, which is more cost-effective than scATAC-seq or sc-multiome, reasoning that TF perturbations that substantially alter the accessibility landscape are also more likely to alter the transcriptome. For each of the 240 candidate TFs, we cloned 3 CRISPRa gRNAs^32^, as well as 75 non-targeting control (NTC) gRNAs, to the CROP-seq vector^33,34^. The resulting 795-guide library was lentivirally packaged and transduced into our CRISPRa mESC line at a low MOI, followed by puromycin selection. After 96 hrs, cells were profiled by scRNA-seq (10x Genomics). 75,340 cells passed quality filtering (median 7,476 UMIs and 2,765 genes per cell), 70% of which were assigned a single gRNA (**Supplementary Fig. 4A-D**). Counting cells with one or more assigned gRNAs, a mean of 240 cells was recovered per target TF. For 192 targets (80%), cells bearing a corresponding gRNA exhibited significantly increased expression of the target TF, by 1.7-fold to 1,916-fold (median 60-fold; **Fig. 1B**; **Supplementary Fig. 4E**). TF upregulation was highly specific, albeit with some exceptions consistent with hierarchical activation (*e.g. Cdx2* → *Hox* genes^35^, *Neurod1* → *Ebf-*family TFs^36^), and activated TFs fell within the expression distribution of natively expressed mESC TFs (**Supplementary Fig. 4F-G**).

We next asked which TFs elicit substantial, coherent changes to the transcriptome. Data visualization highlighted distinct subpopulations, including 2C-like (*Zscan4d+*), fibroblast-like (*Col1a1+*), trophectoderm-like (*Tfap2c+*), trophoblast-like (*Gcm1+*), and pluripotent (*Nanog+*) states, with general agreement between gRNA detection and TF expression (**Fig. 1C-D**). Comparison to NTC-bearing cells identified a median of 10 differentially expressed genes (DEGs) per target TF (IQR, 4-26; range 0-394; **Fig. 1E**; **Supplementary Table 1**), often corresponding to their established roles (*e.g. Sox7* → extraembryonic endoderm (ExEn) markers^37^; *Neurod1* → *Ebf-*family TFs; *Elf5* → placental development) (**Fig. 1F**).

To prioritize target TFs by a quantitative measure of a coherent impact on the transcriptome, we ranked gRNAs by Moran’s I, a test for spatial autocorrelation previously applied to identify variably expressed genes in scRNA-seq data^38^. Many TF-targeting gRNAs showed significant clustering in low-dimensional space, whereas NTC gRNAs did not (**Fig. 1G**). As an orthogonal approach, we also quantified enrichment in local KNN-graph neighborhoods in high-dimensional PCA space (lochNESS score^39^). Reassuringly, 47 of the top 60 TFs were nominated by both metrics (78% overlap; **Fig. 1H**). We therefore moved forward with the top 60 TFs as ranked by the Moran’s I metric to a more highly powered sc-multiome screen.

### A CRISPRa sc-multiome screen of 60 developmental TFs in mESCs

We next sought to measure the impact of these 60 TFs on the chromatin landscape of mESCs through a secondary sc-CRISPRa screen, better powered and read out by a co-assay of both accessibility and gene expression (**Fig. 2A**). After quality control, we recovered 139,374 cell profiles (median 1,914 RNA UMIs and 5,825 unique ATAC fragments per cell; 314,609 peaks called; median peak width = 448 bp; **Supplementary Fig. 5A-B**). At least one gRNA was detected in 58% of cells (median 1,828 cells per TF; **Supplementary Fig. 5C-D**). Activation remained robust, in that gRNA-bearing cells for 56 of 60 (93%) targets showed a significant, ≥2-fold increase in target TF expression, relative to NTC cells, and activation levels correlated well with those from the 240-TF scRNA-seq screen (Pearson’s r = 0.76) (**Supplementary Fig. 5E-F**).

**Figure 2.**
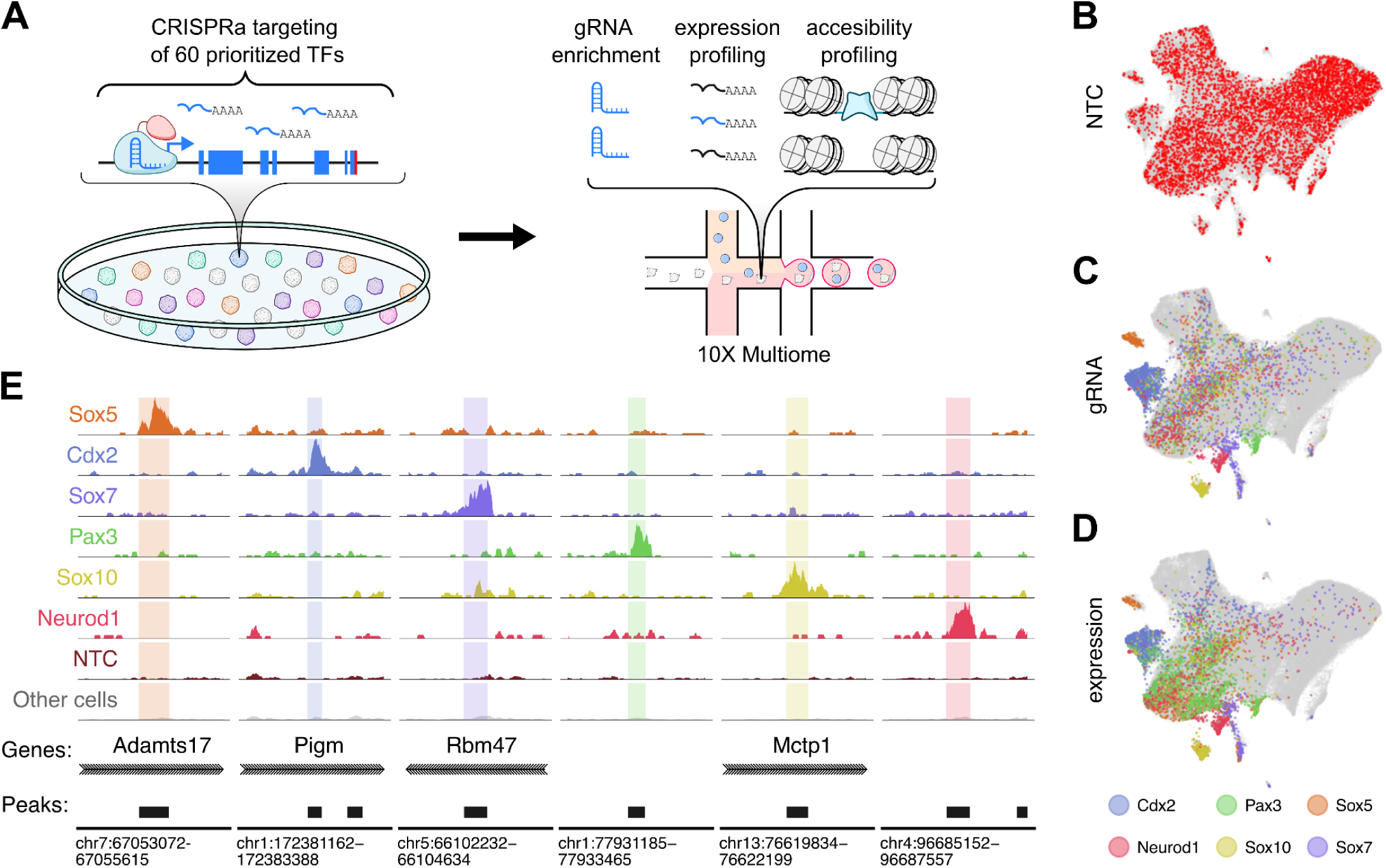
A CRISPRa sc-multiome screen of 60 developmental TFs in mESCs. **(A)** Experimental schematic. Sixty TFs prioritized by the pre-screen were the focus of a new, well-powered CRISPRa sc-multiome screen. **(B)** UMAP embedding of ATAC profiles, highlighting distribution of NTC gRNA(+) cells. **(C)** Same as in **B**, highlighting distribution of lochNESS-filtered gRNA(+) cells targeting *Cdx2*, *Neurod1*, *Pax3*, *Sox5*, *Sox7* and *Sox10*. This filter retains only gRNA(+) cells whose ATAC neighborhood supports successful perturbation (lochNESS > 1). **(D)** Same ATAC UMAP embedding as in **B-C**, highlighting cells in which a given marker TF was both detected and locally enriched among its neighbors (lochNESS > 2). Note that lochNESS is used in distinct ways in **C** (per-cell filtering for high-confidence gRNA(+) detection) and **D** (to enhance visualization). **(E)** Chromatin accessibility read pileups at selected loci bearing DG sites for lochNESS-filtered gRNA(+) cells targeting *Cdx2*, *Neurod1*, *Pax3*, *Sox5*, *Sox7*, and *Sox10*. The bottom two rows show pileups at these same regions for NTC gRNA(+) cells and all unassigned cells.

Because CRISPRa is not uniform across cells, gRNA detection alone is an imperfect proxy for successful perturbation, and cells carrying a TF-targeting gRNA that never experienced activation dilute downstream analyses. Selecting instead on detected target TF expression is also unsatisfactory, both because scRNA-seq sparsity would result in the discarding of actually perturbed cells, and because some targets may be activated downstream of other targeted TFs. We therefore retained, among the cells assigned a gRNA for a given TF, only those whose high-dimensional ATAC profile resembled that of other cells bearing a gRNA for the same TF, as scored by a local cellular heuristic neighborhood enrichment specificity score (lochNESS^39^; **Supplementary Fig. 6A-C**). For clarity, this score is used only to remove gRNA-assigned cells that appear unperturbed, rather than for assigning perturbation identity to cells lacking a detected gRNA. With lochNESS filtering, the UMAP embedding of accessibility profiles revealed focal clusters enriched for gRNAs targeting TFs including *Cdx2*, *Neurod1*, *Pax3*, *Sox5*, *Sox7* and *Sox10*, and for expression of those TFs (**Fig. 2B-D**; **Supplementary Fig. 6D-F**). Many other TFs showed similar but less pronounced clustering, and coverage was sufficient to detect chromatin opening at individual loci relative to NTC cells (**Fig. 2E**).

We then tested for differential accessibility between TF-targeting vs NTC gRNA(+) cells, using bootstrapped pseudo-replicates and an empirical 5% FDR cutoff determined from NTC cells. Across all targeted TFs, 12% of all peaks exhibited differential accessibility (DA), with more examples of sites gained (DG; 28,264) than lost (DL; 10,957) (**Supplementary Table 2**). Per TF, the number of DG sites ranged from 0 (14 TFs) to 14,047 (*Cdx2*) (median 108, mean 871 ± 2,245) and DEGs from 9 to 493 (*Neurod1*) (median 137, mean 145 ± 104). Both the number of DG sites and DEGs increased with power (*i.e.* the number of cells assigned to a TF), but correlated even more strongly with one another, (**Supplementary Fig. 7A-C**). This is a retrospective endorsement of our low-cost transcriptional pre-screen, although we cannot exclude that hypothetical ‘chromatin-openers-only’ are missed. Consistent with their mediating the observed expression changes, peaks proximal to the TSSs of DEGs gained accessibility in targeted cells (**Supplementary Fig. 7D-F**).

*Characterization of TF-specific and shared changes in chromatin accessibility*.

To assess whether DA sites occur at promoters or distal regions, we examined distances to the nearest TSS for non-DA peaks, DG sites or DL sites for any TF, and GC-matched control regions. Non-DA peaks exhibited the expected bimodal distribution, with a promoter-proximal mode but most peaks located distally. DG, DL and control regions were similarly distributed, but with a greatly reduced promoter-proximal mode, consistent with TF-induced accessibility changes primarily occurring at distal enhancers (**Fig. 3A**). On average, DG and DL regions fell closer to TSSs than control regions (median distance 51.6 (DG), 39.5 (DL) vs 57.2 kb (control)).

**Figure 3.**
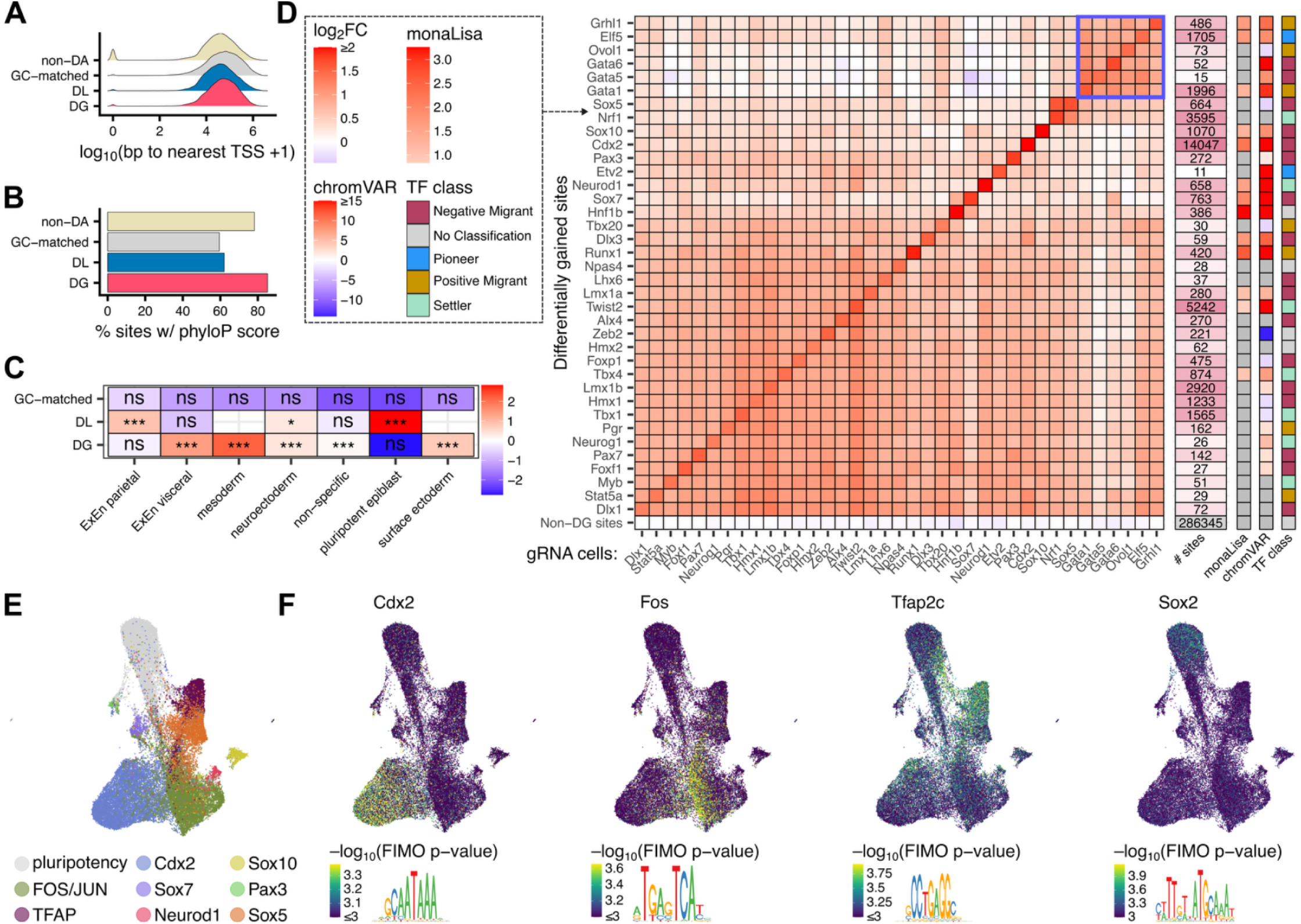
Characterization of TF-specific and shared changes in chromatin accessibility. **(A)** Density plots of distance to nearest TSS for DG, DL, non-DA, and GC- and width-matched background regions. **(B)** % of sites with a phyloP score, for the same groups. **(C)** Heatmap of enrichment for the same groups’ overlap with cell-type-specific accessible peaks from mEBs. Fill indicates fold-enrichment relative to non-DA sites; comparisons with <50 overlapping peaks in either group are excluded. *BH-adjusted p < 0.05, **p < 0.01, ***p < 0.001; ns, not significant. **(D)** Heatmap of average log_2_ fold-change in chromatin accessibility (gRNA vs NTC) at DG sites for each TF (rows) across all other lochNESS-filtered gRNA(+) cell populations (columns), irrespective of significance. Values ≥2 are capped; only TFs with ≥10 DG sites shown; peaks not called DG for any TF are in the bottom row. Adjacent panels, from left to right: number of DG sites per TF; enrichment of matching TF motif family (top enriched motif with −log_10_-adjusted p > 2); chromVAR-derived differences in genome-wide accessibility at that motif family relative to NTC gRNAs (top motif with adjusted p < 0.05); and chromatin-opening classification^45^ of each TF. Blue rectangle highlights block of TFs associated with trophectoderm-like cluster. **(E)** UMAP of DA sites, each point a peak, colored by the peak type to which it was assigned on the basis of its pattern of DG calls across TFs (**Supplementary Fig. 9**). **(F)** As in **E**, colored by the maximum −log₁₀ FIMO p-value motif match (BH-corrected, capped as indicated) for the indicated motifs (*Cdx2*, *Fos*, *Tfap2c*, *Sox2*).

Three additional lines of evidence support the conclusion that TF-induced DG sites are enriched for bona fide, cell type-specific distal enhancers. <u>First</u>, although overall phyloP distributions were broadly similar across peak categories, the fraction of peaks under evolutionary constraint was slightly higher for DG sites (**Fig. 3B**; **Supplementary Fig. 7G-H**). <u>Second</u>, relative to non-DA sites, DG sites overlapped cell type-specific accessible sites from all three germ layers of mouse embryoid bodies (mEBs) significantly more often, whereas DL sites were instead enriched for overlap with pluripotent-like mEB cells^40,41^ (**Fig. 3C**). <u>Third</u>, DG sites carried little evidence of activity in an ENCODE mESC cell line (E14), as only 15% overlapped ChIP-seq-ascertained histone modifications^42–44^, versus 32% of non-DA, 55% of DL and 11% of control regions, while overlap with the enhancer marks H3K4me1 and H3K27ac was far lower for DG than DL sites (DG: 11% and 1.4%; DL: 43% and 24%) (**Supplementary Fig. 7I-K**). DL sites thus appear to correspond to distal, pluripotency-specific elements that lose accessibility upon differentiation, whereas DG regions appear largely inactive in wild-type mESCs. Overlap with H3K9me3 was minimal throughout (<1%), consistent with chromatin-opening TFs rarely accessing constitutive heterochromatin.

To assess which DG sites were TF-specific, we calculated the mean accessibility fold-change in each TF’s DG regions for cells assigned to other TFs, revealing a continuum of specificity (**Fig. 3D**). DG sites for some TFs were highly specific (*e.g. Cdx2*, *Neurod1*, *Sox10*); others were shared among a limited subset (*e.g. Elf5*, *Grhl1*, *Gata1*, *Gata5*, *Gata6*, *Ovol1*); and still others increased across most TFs, possibly reflecting differentiation modules shared across fates. For 13 TFs, the cognate motif was significantly enriched in DG sites, consistent with primary rather than secondary effects. Only two of the 37 TFs with ≥10 DG sites, *Elf5* and *Etv2*, had been computationally classified as pioneer TFs^45,46^, although pioneering activity has been reported in specific contexts for several others (see **Discussion**). Applying similar analyses to DL sites, the TFs with the most DL sites were those with the most DG sites, and cognate motif enrichment was limited to *Sox5*, *Sox7* and, weakly, *Elf5* (**Supplementary Fig. 8**). Because the *Sox5* and *Sox7* motifs closely resemble that of *Sox2*, we suspect that most DL sites reflect secondary loss of SOX2-occupied sites rather than sequence-specific repression.

As an orthogonal view on these data, we performed dimensionality reduction on the transposition of the cell × peak matrix^47^, where peaks with shared accessibility across cells will group together. Annotating peaks by the TFs for which they were called DG yielded nine major “peak types”, which we then characterized by motif enrichment (**Fig. 3E-F**; **Supplementary Fig. 9**). These included: (i) six types comprising sites called DG exclusively for *Cdx2*, *Sox7*, *Neurod1*, *Sox10*, *Pax3* or *Sox5* (for *Sox5*, sites DG for *Sox5* and/or *Nrf1,* which share a gRNA with sequence similarity; **Supplementary Fig. 6G-H**), enriched for the cognate motif in the first four cases and for DMRT1 and GATA motifs in the cases of *Pax3* and *Sox5*, respectively; (ii) one type comprising sites DG for one or more of *Elf5*, *Grhl1*, *Gata1*, *Gata5*, *Gata6* and *Ovol1* and no other TF, enriched for TFAP2A-E motifs (“TFAP”); (iii) one type comprising the remaining DG sites, with FOS/JUN among its top motifs (“FOS/JUN”); and (iv) one type comprising sites called DL for any TF and DG for none, enriched for SOX2 and POU5F1 motifs (“pluripotency”).

Because DA testing is limited by scATAC-seq data sparsity, we performed a complementary analysis at the motif level with chromVAR^48^. Pseudobulking lochNESS-filtered cells by individual gRNA, so that independent gRNAs targeting the same TF served as replicates, we asked whether genomic occurrences of the TF’s motif gained accessibility relative to NTC cells, independent of peak calls. Twenty-two TFs showed significant genome-wide gains in cognate motif accessibility (p-adj. < 0.05), in good agreement with motif enrichment in DG sites (**Fig. 3D**). Some signals were not strictly TF-specific, *e.g.* NFE2 motif accessibility increased in association with multiple target TFs, suggesting shared and likely secondary effects (**Supplementary Fig. 10**).

Together, these results showcase striking heterogeneity in the capacity of newly expressed TFs to reshape chromatin, ranging from little detectable remodeling to the opening of thousands of previously inactive sites. Responses varied in the number and specificity of sites gained and their concordance with the TF’s cognate motif, with four broad patterns. <u>First</u>, some TFs drove strong transcriptional responses with little to no detectable chromatin remodeling (*e.g. Npas4*, *Alx4*). <u>Second</u>, some TFs drove highly specific gains with strong enrichment of their cognate motif (*e.g. Cdx2*, *Neurod1*, *Sox10*), consistent with direct chromatin opening. <u>Third</u>, some drove similarly specific gains but with little enrichment of their cognate motif, instead showing strong enrichment for other TFs’ motifs (*e.g.* DMRT motifs for *Pax3*), suggesting much of their remodeling occurs indirectly through downstream TFs. <u>Fourth</u>, some TFs with distinct motif preferences converged on shared accessibility programs, including site-groups associated with TFAP or FOS/JUN motifs. Thus, in many cases, the identity of the activated TF alone is insufficient to explain which sites become accessible.

### Hierarchical co-TF usage in the ExEn-like state

Differentially gained sites shared across targeted TFs with different motif preferences could reflect common use of a co-TF already expressed in mESCs (“opportunistic”), downstream activation of a common co-TF (“hierarchical”), or secondary differentiation effects that lead to shared accessibility changes. As a case study in how to distinguish between these scenarios, we focused on a group of cells, defined by Louvain clustering on ATAC data, with extraembryonic endoderm-like (ExEn) expression^37^. This cluster was enriched for *Sox7*, *Gata6* and *Hnf1b* gRNA(+) cells (**Fig. 4A**), while DG sites associated with this cluster were strongly enriched for SOX, GATA and HNF1B motifs (**Supplementary Fig. 11A, E, G-H**).

**Figure 4.**
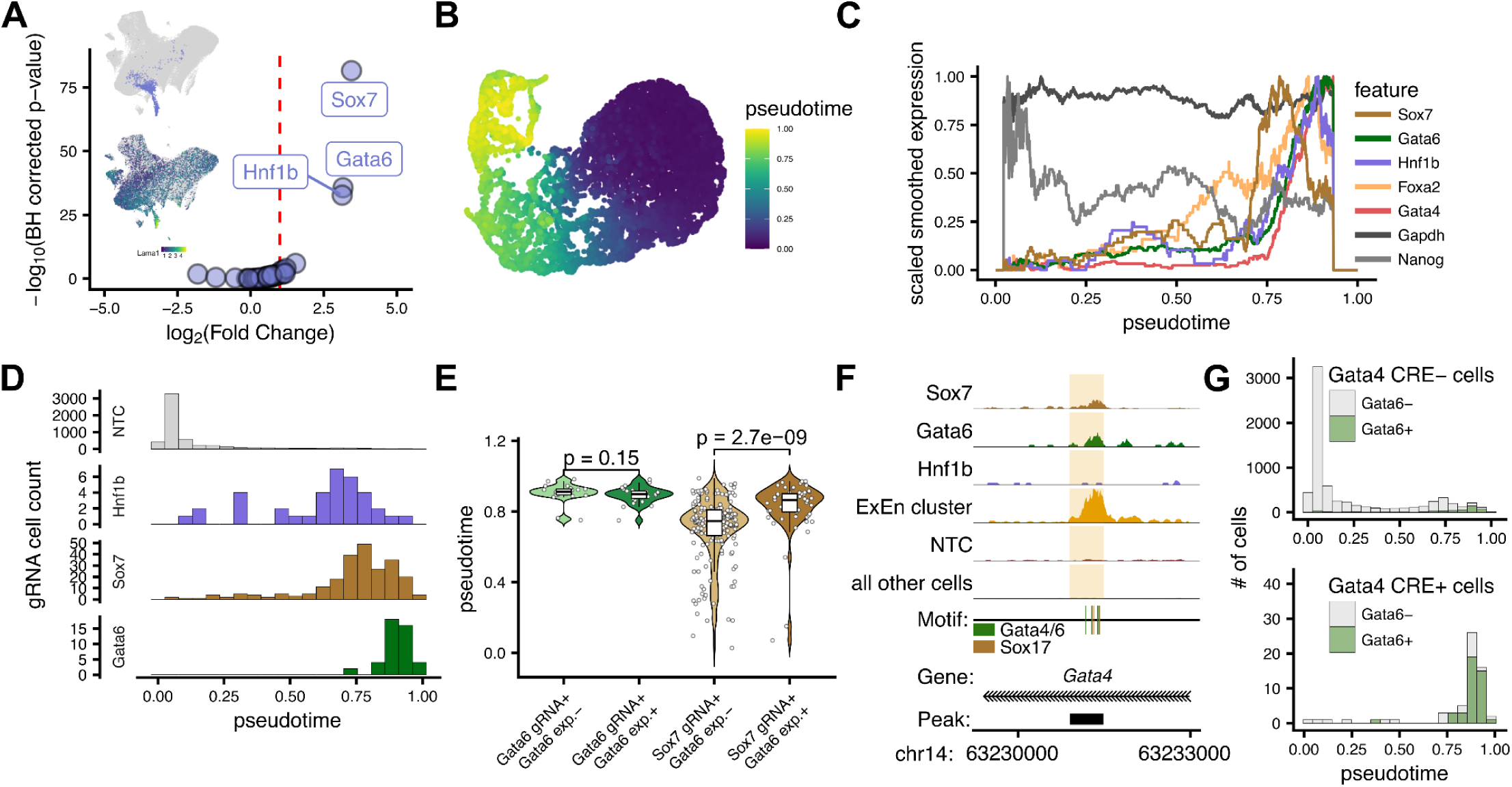
Hierarchical co-TF usage in the ExEn-like state. **(A)** Volcano plot showing enrichment of gRNA(+) cells for each target TF within the ExEn cluster. The x-axis shows log_2_-fold difference in proportion of target gRNA(+) vs NTC cells assigned to the cluster. The y-axis shows −log_10_(FDR). The inset ATAC UMAPs highlight the ExEn cluster (top) and expression of ExEn-marker *Lama1*^52^ (bottom). **(B)** UMAP visualization of a diffusion pseudotime embedding constructed from all cells from the ExEn-like cluster together with NTC gRNA(+) cells. Pseudotime values inferred from the diffusion map are overlaid onto the embedding. **(C)** Smoothed expression profiles across pseudotime for the targeted TFs *Gata6*, *Sox7* and *Hnf1b*, together with a representative pluripotency marker (*Nanog*), a housekeeping gene (*Gapdh*), a TF involved in ExEn formation (*Foxa2*)^53^, and an ExEn master regulator (*Gata4*)^49^. **(D)** Histograms of pseudotime values for NTC, *Hnf1b*, *Sox7*, and *Gata6* gRNA(+) cells from the ExEn trajectory. Each facet displays the pseudotime distribution for cells assigned the indicated gRNA class, with independent y-axis scaling for visualization. **(E)** Violin plots showing pseudotime distributions for *Gata6* gRNA(+) and *Sox7*-gRNA(+) cells in the ExEn trajectory, further stratified by detectable *Gata6* expression (*Gata6* exp.+, ≥1 UMI). Points represent individual cells, and overlaid boxplots indicate the median and interquartile range. P-values from two-sided Wilcoxon rank-sum test. **(F)** ATAC read pileup at a highly conserved, intronic *Gata4* CRE^51^ in *Sox7* gRNA(+) cells, *Gata6* gRNA(+) cells, *Hnf1b* gRNA(+) cells, ExEn-annotated cells, NTC cells, and all other cells. **(G)** Histograms of pseudotime values for cells without (top) or with (bottom) detected accessibility at the intronic *Gata4* CRE. In each bar, cells are further stratified by *Gata6* expression.

Examining the cross-regulation of these three TFs and other ExEn regulators, we found that both *Sox7* and *Gata6* gRNA(+) cells upregulate the ExEn master regulator *Gata4*^49^; while *Sox7* gRNA(+) cells also activate *Gata6*, but not vice versa (**Supplementary Fig. 12A-C**). Consistent with these relationships, both *Sox7* and *Gata6* gRNA(+) cells exhibited increased accessibility at GATA6 motifs, whereas increased accessibility at Sox7 motifs was restricted to *Sox7* gRNA(+) cells (**Supplementary Fig. 12E**).

We asked whether these putative hierarchical interactions are reflected in a pseudotime trajectory, constructed from genes and sites differentially expressed or accessible between ExEn regulator gRNA(+) vs NTC gRNA(+) cells^12^ (**Fig. 4B**; **Supplementary Fig. 13A**). While housekeeping genes remained consistently expressed across pseudotime, pluripotency-associated genes decreased and ExEn genes progressively increased (**Fig. 4C**; **Supplementary Fig. 13B**). Furthermore, cells assigned a targeting gRNA were distributed along progressively later regions of the trajectory, orienting the pseudotime with the progression of differentiation (**Supplementary Fig. 13C**). *Hnf1b* gRNA(+) cells were earliest, consistent with a more limited induction of the ExEn program (*e.g.* a lack of *Gata4* expression), followed by *Sox7* and finally *Gata6* gRNA(+) cells (**Fig. 4D**). *Sox7* expression rose earliest, with *Gata6* peaking later (**Fig. 4C; Supplementary Fig. 13D-F**).

Consistent with a hierarchical model in which *Sox7* activates *Gata6*, *Sox7* gRNA(+) cells with detectable endogenous *Gata6* expression were shifted toward later pseudotimes than *Sox7* gRNA(+) cells lacking detectable *Gata6* transcripts, whereas *Gata6* gRNA(+) cells showed no analogous difference (**Fig. 4E**), possibly because even though *Gata6* CRISPRa activation is robust (**Supplementary Fig. 12A-D**), dropout nonetheless limits the detection of *Gata6* transcripts in some *Gata6* gRNA(+) cells.

Extending this framework to chromatin accessibility also revealed distinct pseudotemporal dynamics across CREs. For example, *Bicc1*, a previously reported downstream target of *Hnf1b* with an intronic CRE bound by *Hnf1b*^50^, increases steadily along the trajectory (**Supplementary Fig. 13G**). Whereas the *Bicc1* promoter was already accessible early in pseudotime, the intronic CRE became accessible predominantly at the latest stage, coinciding with maximal *Hnf1b* and *Bicc1* expression and promoter accessibility (**Supplementary Fig. 13G**).

Given the induction of *Gata4* by *Sox7* and *Gata6*, but not *Hnf1b*, we also examined a highly conserved intronic CRE residing 40 kb downstream of the *Gata4* TSS that bears numerous SOX and GATA motifs^51^. This element showed increased accessibility in ExEn-like cells, including *Sox7*- and *Gata6*-gRNA(+) cells, but not *Hnf1b*-gRNA(+) cells (**Fig. 4F**). Cells with accessibility at this CRE were enriched toward later pseudotime and strongly enriched for *Gata6* expression (**Fig. 4G**), consistent with both features marking the most advanced ExEn-like states. Together, these findings illustrate how, even in the context of a CRISPRa screen harvested at a single timepoint, joint RNA and chromatin accessibility measurements can nominate potential temporal orderings of regulatory events during TF-induced cell state transitions.

### Opportunistic co-TF usage of heterogeneously expressed Tfap2c

Interestingly, *Gata6* gRNA(+) cells were enriched not only in the cluster with ExEn-like expression (**Fig. 4A**), but also in a second cluster with trophectoderm-like expression (**Fig. 5A**), sharing DG sites with *Elf5*, *Gata1*, *Gata5*, *Grhl1* and *Ovol1* gRNA(+) cells (**Fig. 3D**). Motif enrichment at these sites revealed the cognate target TF motifs, but also enrichment of the TFAP2C motif, whose cognate TF, *Tfap2c*, is specifically expressed in this cluster (**Fig. 5B-C**; **Supplementary Fig. 9**; **Supplementary Fig. 11B-F,I**).

**Figure 5.**
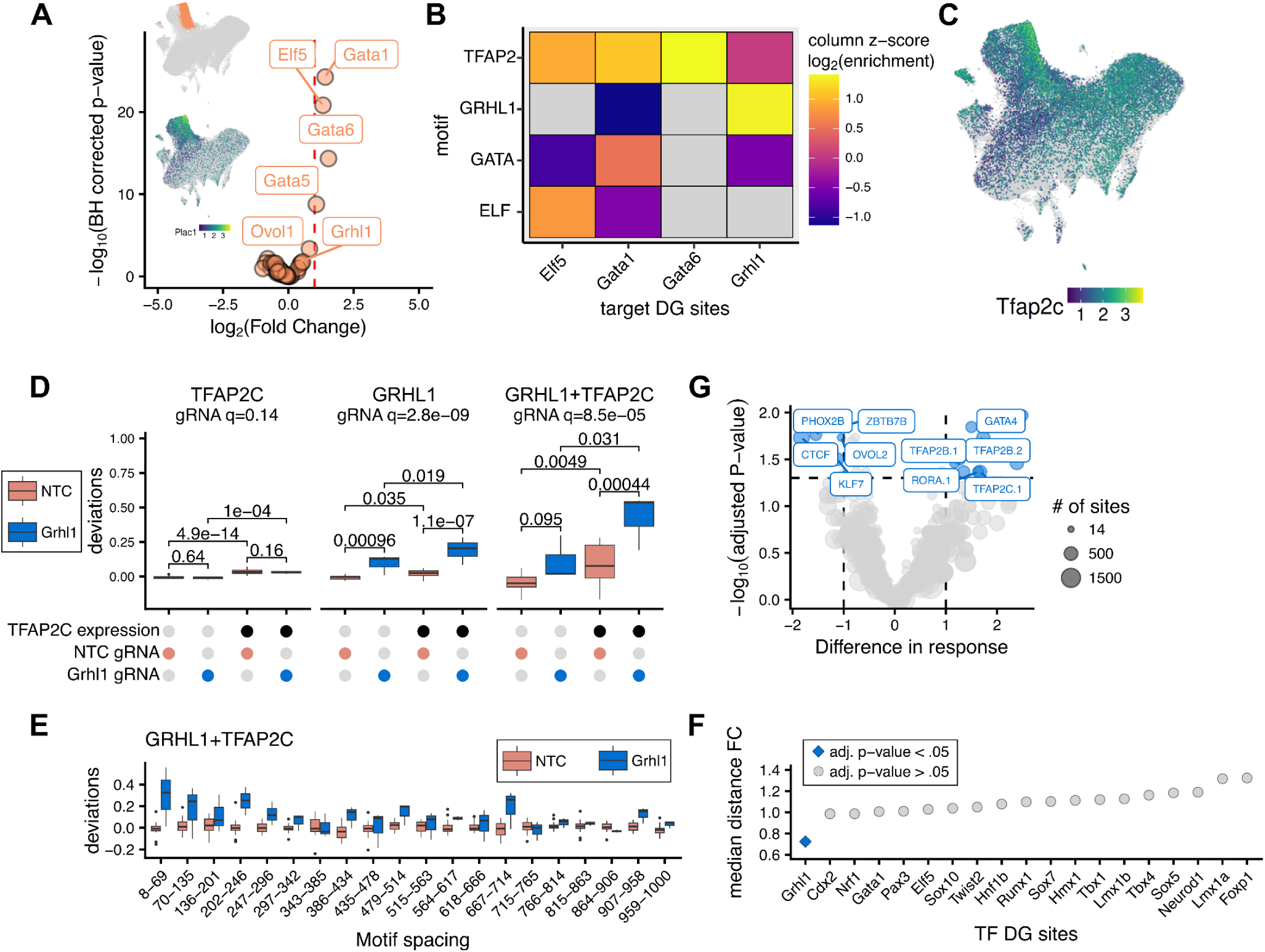
Opportunistic co-TF usage of heterogeneously expressed *Tfap2c*. **(A)** Volcano plot of enrichment of each gRNA within the trophectoderm-like cluster; x-axis, log_2_ fold-difference in the proportion of target cells vs proportion of NTC cells. y-axis, −log_10_(Benjamini-Hochberg-corrected p-value). Top inset, ATAC UMAP colored by cluster; bottom inset, expression distribution of the trophoblast marker *Plac1* in the ATAC UMAP. **(B)** Heatmap of motif enrichment (via monaLisa) for each motif family (rows) and each TF’s DG sites (columns). For each TF-motif-family pair, fill shows the mean log_2_(enrichment) across significant motifs in that family, z-score normalized within each column; tiles with no significant motif in the family are grey. **(C)** Expression distribution of *Tfap2c* in the ATAC UMAP embedding. **(D)** Boxplots of chromVAR deviations for the TFAP2C motif, GRHL1 motif, and GRHL1-TFAP2C co-motif instances after stratifying and pseudobulking *Grhl1* and NTC cells by detection of *Tfap2c* expression. *Grhl1* is shown as an exemplar, with other TFs in **Supplementary Fig. 16**. Adjusted p-values for between-group comparisons are shown. Above each motif class, the significance of the marginal CRISPRa effect averaged across TFAP2C strata is indicated (“gRNA q =”). **(E)** chromVAR deviation scores for genomic regions containing the GRHL1-TFAP2C co-motif, binned by inter-motif distance (x-axis, with bin ranges indicated). Boxplots show the distribution of deviation scores across pseudobulked *Grhl1* and NTC gRNA(+) cells. (**F**) Dot plot of the fold change in median GRHL1-TFAP2C motif distance between each TF’s DG peaks and all other peaks containing both motifs (median distance in DG sites / median distance in other sites). Only TFs with >50 DG peaks were included. TFs are ordered by FC, and point shape indicates statistical significance after Benjamini-Hochberg multiple testing correction of Wilcoxon p-values. (**G**) Volcano plot showing the difference in chromVAR accessibility response^55^ between sites bearing a GRHL1 motif together with the indicated co-motif, versus sites bearing a GRHL1 motif alone; x-axis, difference in response; y-axis, −log_10_(adjusted p-value). Motifs in blue had an absolute difference ≥1 (adjusted p-value < 0.05); the top five significant motifs with the largest effect sizes in each direction are labeled.

What might explain this, given that we did not target *Tfap2c* by CRISPRa? One possibility is that the association is hierarchical, *i.e.* that these targeted TFs converge on the direct or indirect activation of *Tfap2c*, explaining their shared accessibility gains. However, relative to UMI-matched NTC cells, *Tfap2c* transcript levels were not elevated, or at most weakly so (max FC = 1.27 among significant comparisons; **Supplementary Fig. 12A**). The same conclusion held under a more stringent test, in which we applied a nearest-neighbor differential expression framework to all target TFs, matching each gRNA(+) cell to a gRNA(−) cell occupying the closest position in ATAC PCA space in order to control for heterogeneity in pre-existing cell state with our alternate modality (**Supplementary Fig. 14A**; **Methods**). In this framework, *Tfap2c* was not called as differentially expressed for any of the perturbation groups enriched in the trophectoderm-like cluster, whereas the targeted TF itself was in most cases called as differentially expressed, along with known or plausible direct regulatory targets (**Supplementary Fig. 14B-D**).

This led us to instead favor an alternative hypothesis, which is that the two outcomes observed for *Gata6* gRNA(+) cells are a consequence *Gata6*’s interaction with “pre-existing” heterogeneity of *Tfap2c* in mESCs, whose expression is 14-fold higher in the trophectoderm-like than in the ExEn-like cluster (**Fig. 4A**; **Fig. 5A,C**; **Supplementary Fig. 12F**) and whose detectable presence in unperturbed mESC was the reason it was excluded from the target TF panel. In this model, CRISPRa-mediated activation of *Gata6* can lead to different outcomes depending on the pre-existing state of the mESC in which the perturbation occurs, with *Tfap2c*-high mESCs adopting the trophectoderm-like fate, and *Tfap2c*-low mESCs adopting the ExEn fate. A further assertion of this model is that the other cluster-associated TFs (*Elf5*, *Gata1*, *Gata5*, *Grhl1*, *Ovol1*) also require a *Tfap2c*-high background to adopt the trophectoderm-like fate, but unlike *Gata6*, do not strongly promote an ExEn-like fate if their activation is superimposed on a *Tfap2c*-low background.

If correct, the association of *Tfap2c* with a trophectoderm-like outcome could be due to *Tfap2c* driving newly accessible sites independently of those driven by the activated TFs. Alternatively, *Tfap2c* and the activated TFs might act cooperatively to open chromatin at sites where their motifs co-occur. A prediction of the latter possibility is that co-occurrences of the motifs of TFAP2C and trophectoderm-associated target TFs should be enriched at DG sites. To test this, we repeated the chromVAR analysis on gRNA(+) cells for these TFs while explicitly including composite motifs falling within 5-100 bp. Pseudobulked *Gata5*, *Gata6*, *Elf5* and *Grhl1* gRNA(+) cells exhibited significant increases in accessibility at their composite motifs with TFAP2C, while *Gata1* showed a non-significant trend (**Supplementary Fig. 15**), consistent with cooperative opening.

To more directly test whether accessibility at these composite sites depends on endogenous *Tfap2c* activity, we stratified gRNA-perturbed cells into *Tfap2c*(+) and *Tfap2c*(-) subsets prior to pseudobulking and quantifying motif deviations (**Fig. 5D**; **Supplementary Fig. 16**). *Tfap2c*(+) cells that were also *Grhl1*, *Elf5* or *Gata6* gRNA(+) tended to exhibit higher accessibility at composite sites than all other groups, providing further support for a model in which *Tfap2c* enhances accessibility co-operatively. By comparison, this was not the case for *Tfap2c*(+), *Sox7* gRNA(+) cells, as expected given that *Sox7* is not associated with the trophectoderm-like cluster. Interestingly, *Tfap2c*(+) NTC gRNA(+) cells displayed elevated accessibility at these sites relative to *Tfap2c*(-) NTC gRNA(+) cells, likely attenuating the interaction signal but also suggesting that *Tfap2c* expression may be sufficient to ‘prime’ these regions for opening prior to the expression of target TFs.

We next asked whether these accessibility changes depend on the distance between the two motifs. Across multiple target TF-TFAP2C pairs, sites harboring heterotypic motifs in closest proximity (<70 bp) exhibited larger increases in accessibility than those with greater inter-motif distances, particularly for *Grhl1* and *Gata6*, whereas *Gata1* showed the largest increase at slightly larger distances (**Fig. 5E**; **Supplementary Fig. 17A-F**). Even though median DG peak size is 535 bp, accessibility neared baseline at distances around 200 bp, in line with cooperativity of pioneers with other TFs’ motifs within nucleosome range, as recently described in unperturbed mESCs^54^. In several cases, we also observed periodically increased accessibility at distances that might represent increased TF occupancy in linker regions of positioned nucleosomes (*e.g.* for GRHL1-TFAP2C at ∼220 and ∼410 bp). Examining GRHL1-TFAP2C motif spacing within *Grhl1* DG sites, sites containing both motifs exhibited a significant ∼30% reduction in spacing (median ∼102 bp) relative to all other peaks containing both motifs (median ∼143 bp), an effect not observed for GRHL1-TFAP2C motif instances in the DG sites of other TFs (**Fig. 5F**; **Supplementary Fig. 17G-I**). When assessing changes in co-motif accessibility across all possible partner motifs^55^ for *Grhl1*, TFAP2C emerged as one of the top significant partners with gained accessibility. Interestingly, this unbiased approach also uncovered motifs whose co-occurence with GRHL1 accompanies a loss of accessibility, *e.g.* the cognate motifs of repressors ZBTB7B, *Hes7* and *Klf7*, as well as the insulator/looping factor CTCF (**Fig. 5G**). Similar putative TF motif interactions were also nominated from multiome Perturb-seq data in a recent study in RPE-1 cells ^8^.

Together, these results support opportunistic co-TF usage as a general feature of this group of TFs. *Elf5*, *Gata6* and *Grhl1* all tend to gain accessibility preferentially in cells that already express *Tfap2c* and at sites where their own motif co-occurs with a TFAP2C motif. *Gata6* additionally illustrates how such dependencies shape which fates are reachable. *Gata6* is the only one of these TFs enriched in both clusters, and its outcome appears to be contingent on the starting state. Under our proposed model, it accesses the shared trophectoderm-like program in *Tfap2c*-high cells, whereas in *Tfap2c*-low cells it opens only a subset of DG sites associated with the ExEn fate. The latter are in turn partly shared with *Sox7* gRNA(+) cells, since *Sox7* activation appears to induce *Gata6*. These TFs thus remain capable of opening sites individually, but reach their fullest accessibility gains, and a distinct set of fates, only where a co-TF is present.

## DISCUSSION

In this CRISPRa sc-multiome screen, targeting 60 developmental TFs and profiling ∼140,000 single cells, we observe a continuum of responses, from no detectable remodeling to the opening of thousands of previously inactive distal elements. Some TFs appear to act autonomously, opening sites enriched for their own motif (“autonomous”). Others depend on a co-TF, either one already heterogeneously expressed among mESCs (“opportunistic”) or one that they themselves induce (“hierarchical”).

Several TFs causing specific accessibility gains in our study have strong literature support as pioneer factors: *Grhl1*^56^; *Gata6*^57^; *Etv2*, shown to bind nucleosomal DNA and recruit *BRG1*^58^; the proneural factors *Neurod1*^59^ and *Neurog2*^23^; and *Pax3*, recently identified as one of the strongest nucleosome-binding pioneer factors^2^. Evidence is conflicting or strongly context-dependent for *Cdx2*, the factor with the most gained sites in our study. It behaves as a pioneer-like master factor in some contexts^60^, but not in the developing gut, where it fails to instruct major shifts in chromatin^61^. Likewise, *Runx1*, which here shows highly specific DG sites alongside motif and chromVAR enrichment, was previously classified as a non-pioneer that only binds accessible chromatin upon ectopic expression in fibroblasts^2^. *Twist2*, which yielded the second most gained sites after *Cdx2*, is a predominantly repressive bHLH factor previously associated with reduced rather than increased accessibility across diverse contexts^62–64^, whereas here we observe the opposite. Our approach is not inherently biased toward detecting opening over closing, yet the only factor showing cognate motif enrichment in a substantial fraction of DL sites, without resembling pluripotency factor motifs, is *Elf5*, which also shows motif enrichment in DG sites and has a previously reported dual role in activation and repression^65^.

Overall, our findings suggest that the ability of a TF to “pioneer” chromatin opening is highly cell-state-specific, nuanced, and continuous, rather than a binary property. Particular co-TFs or chromatin states may be required for a TF to exert chromatin-remodeling capacity, and may even determine whether it opens or closes chromatin. For example, *Grhl1*, an established pioneer TF, opens chromatin in our study, but our data suggest this occurs mainly in the presence of *Tfap2c* in the same cell, a dependency that to our knowledge has not been previously described (though a GRHL2-TFAP2A motif interaction has been^66^). This example also illustrates how single-cell, single-TF, pooled perturbation screens, despite reduced signal-to-noise relative to bulk assays, can exploit cell state heterogeneity present within even a single cell type to discover such interactions.

The two dependent classes reach similar endpoints by different routes. *Sox7* induces *Gata6* and then acts alongside it, such that the co-TF is a consequence of the perturbation (“hierarchical”). *Grhl1* and several other TFs instead exploit *Tfap2c* where it is already expressed (“opportunistic”). *Gata6* illustrates the consequences most directly. In *Tfap2c*-high cells it accesses the shared trophectoderm-like program, whereas in *Tfap2c*-low cells it opens only the subset of sites associated with the ExEn fate. The same perturbation applied to two cell states therefore appears to yield two outcomes, a reminder that which fates are reachable from a given starting point is also a function of the cell state a TF lands in, not of the TF alone.

Start state heterogeneity is thus informative, but also confounding, and it is difficult to disentangle from primary and secondary perturbation effects in a single snapshot. Other strategies for disentangling pre-existing heterogeneity, direct and secondary regulatory effects, might include a multiome time-series, or perhaps recently described “molecular memory” methods such as TimeVault or Recordanelles^67,68^.

A growing corpus of steady-state, genome-wide chromatin accessibility profiles, generated across cell types and organisms, is enabling computational TF binding site prediction at a scale out of reach for per-TF ChIP-seq. These predictions are accurate for TFs that bind similar sites across cell states, such as CTCF, but overall AUPRC still hovers below 0.3 for held-out TF-context combinations, and is even worse for TFs with high cell state specificity, exactly the group for which predictions would be most valuable^69^. Furthermore, predicting gene expression from TF binding remains unsolved^70^. Both challenges are more pronounced outside steady-state, *i.e.* when asking how a TF will act when newly upregulated or downregulated in a given cell type. Systematic TF perturbations at scale, read out by multiome assays of gene expression and chromatin accessibility, promise to advance our ability to address these challenges, *e.g.* by identifying the regulatory regions and genes whose activities change as a direct or indirect consequence of a specific perturbation. Chromatin accessibility is particularly well suited to this, reporting on the regulatory interactions upstream of gene expression changes and providing a direct link to DNA sequence; models trained on observational ATAC or multiome data already show promise in predicting perturbation effects, often outperforming models that take purely transcriptomic input^71–75^. No models are yet trained on multiomic perturbation data, but further datasets, including the one reported here, may motivate building them, helping to fill the gap between DNA language- and transcriptome-based single-cell foundation models. Even our simple correlation-based analyses nominate regulatory interactions between TFs that would be challenging to detect from RNA alone.

There are several limitations to our study. <u>First</u>, differential activation efficiency and data sparsity make it challenging to quantify relative pioneering capacity, or to definitively conclude that TFs for which we do not observe strong changes lack chromatin-opening capability. <u>Second</u>, our assignment of sites to TFs rests on changes in accessibility and motif enrichment rather than direct measurement of binding, and since most TFs harbor their cognate motif in fewer than 25% of their ChIP-seq binding sites^69^ distinguishing directly bound sites from downstream effects will require orthogonal approaches. <u>Third</u>, we activate TFs against the background of only a single starting cell type, mESCs. Some of our observations may be unique to stem cells, not only because of the context-specificity of TF activity but also because of the chromatin features that come with pluripotency.

Despite these limitations, we anticipate that this framework, especially if coupled with improvements to the scalability of sc-multiome screens^76^, could be applied to additional starting cell states to systematically map how the ability of TFs to remodel cell state varies across contexts, as well as mechanisms underlying any dependencies. Because single-TF perturbations in a heterogeneous starting population can nominate TF-TF interactions, such screens could also identify candidate pairs on which to focus higher-order perturbation screens, within a combinatorial space that may be impractically large to tackle in an unsupervised manner.

Over the past decade, the manifold of mammalian cell states has been extensively well-charted, but almost exclusively through single-cell measurements of gene expression. TF perturbation screens that also rely purely on gene expression may inform how to move around the manifold, but offer little insight into what underlies these movements, nor how to navigate the manifold with intention. Our results, based on a TF perturbation screen in which both chromatin and gene expression were co-assayed, suggest that a TF’s ability to change cell state is highly dependent on both the *cis-* and *trans-* context in which it finds itself, with some TFs acting autonomously, some opportunistically, on the back of co-TFs already present, and some hierarchically, by inducing their own partners. These behaviors play out against a background of enhancers whose motif content imposes combinatorial requirements, so that only certain combinations of TFs open certain sites, and the paths available to a cell are those the genomic landscape allows. Looking forward, we anticipate that charting such *cis-* and *trans-* dependencies will result in a more satisfying map of the mammalian cell state manifold, one that advances our understanding of endogenous gene regulation in development and beyond, while also enabling its intentional navigation.

## AUTHOR INFORMATION

## Supporting information

TablesS1-3

## Acknowledgments

We are grateful to members of the Shendure Lab and William Stafford Noble for helpful discussions and suggestions on experimentation, data analysis, and visualization. This work was supported by the NIH (R01HG010632 to J.S.), Paul G. Allen Frontiers Group (Allen Discovery Center for Cell Lineage Tracing to J.S.), Seattle Hub for Synthetic Biology, a collaboration between the Allen Institute, Chan Zuckerberg Initiative and University of Washington (award number CZIF2023-008738 to J.S.), Swiss National Science Foundation (TMSGI3_211400 to S.D.), and Brotman Baty Institute for Precision Medicine (J.S.). J.S. is an Investigator of the Howard Hughes Medical Institute.

## Author Contributions

S.D. and J.S. initiated, conceptualized, and supervised the study. A.K. and S.D. performed experiments with help from R.D. and S.J.. A.K. performed the data analyses with guidance and input from S.D. and help from C.Q.. P.L.G. developed the epiwraps and chromVAR-based co-motif analysis approach, which was implemented and adapted by A.K.. S.D., A.K., and J.S. wrote the manuscript with input from all authors.

## Data & Code Availability

Raw sequencing data and associated processed data have been deposited in the Gene Expression Omnibus (GEO) under accession number GSE353547. Code used for analyses and figures have been deposited on GitHub at https://github.com/shendurelab/chromatin-opening-TFs

## Supplementary Tables

Supplementary Table 1. Differentially expressed genes for each targeted TF (relative to NTC cells) for the 240-TF scRNA-seq screen.

Supplementary Table 2. Differentially accessible sites for each targeted TF (relative to NTC cells) for the 60-TF Multiome screen.

Supplementary Table 3. All spacer sequences used in the study.

## METHODS

### Experimental methods

#### Mouse embryonic stem cell culture

Mouse embryonic stem cells were maintained in N2B27 medium supplemented with inhibitors (2i medium). N2B27 medium is composed of 47.5% DMEM/F12 medium with HEPES (Thermo Fisher, cat#11330032), 47.5% Neurobasal medium (Thermo Fisher, cat#21103049), 1% N2 Supplement (Thermo Fisher, cat#17502048), 2% B27 supplement (Thermo Fisher, cat#17504044), 1% Glutamax (Thermo Fisher, cat#35050061), 1% MEM non-essential amino acids (Thermo Fisher, cat#11140050), 0.001% β-mercaptoethanol (Thermo Fisher, cat#21985023). Before each use, 5uL of 10mM MEK inhibitor (Selleck Chemicals cat#S1036) and 15uL of 10 mM GSK inhibitor (Selleck Chemicals cat#S2924) were added to 50mL of N2B27 media. Cells were cultured on plates that were pre-treated with 0.2% gelatin solution, and passaged every 2 days. For routine passaging, cells were washed using DMEM/F12 (Thermo Fisher, cat#11330032) supplemented with 0.03% BSA (Sigma cat#9048-46-8) and dissociated from culture vessels using Accutase (Thermo Fisher, cat#A1110501) to generate single-cell suspensions prior to replating.

#### Generation of monoclonal CRISPRa cell line

Rex1-GFPd2 mouse embryonic stem cells (a kind gift from the Austin Smith lab^77^) – in which the coding sequence of one allele of Rex1 (gene name *Zfp42*) has been replaced with a GFPd2-IRES-bsd cassette that produces a destabilised version of GFP protein as a reporter for naïve pluripotency – were used to generate a monoclonal CRISPRa line. SP-dCas9 with VP64-p65-Rta (VPR) fused to the C-terminus (Addgene #63798)^78^ was cloned into a PiggyBac transposon backbone (PB-EF1α-MCS-IRES-RFP, Systems Biosciences #PB531A-2) under control of an EF1alpha promoter. The resulting vector was transfected into the Rex1-GFPd2 mESC together with transposase plasmid (Systems Biosciences #PB210PA-1) using standard protocols. RFP-positive cells were sorted by FACS and then seeded at low density to grow monoclonal lines. Several individual clones were picked and their CRISPRa activity was tested by transient transfection with gRNAs targeting transcription factors not expressed in mESC, followed by qRT-PCR (*Ascl1* gRNA GGGCGCGGGCAACTGGAGGG from Horlbeck et al.^32^; for cloning into CROP-seq vector and qRT-PCR see below). The clone with the highest activation levels was selected for the downstream experiments.

#### Selection of 240 murine TFs to screen and their corresponding spacer sequences

Candidate TFs were based on the TOME (Transcriptional Ontogeny of Mammalian Embryogenesis) dataset^31^, with the aim of selecting for TFs that were associated with the emergence of one or more cell types in early development. Based on cell-state relationships inferred on the basis of transcriptional similarity, 531 TF were identified that met all of the following criteria: (i) significantly upregulated in a newly emerged cell type, relative to the pseudoancestor; (ii) detected in at least 10% of cells in a newly emerged cell type; and (iii) not significantly upregulated at any ‘sister’ edges, relative to a newly emerged cell type^31^. TFs were further filtered to only those with an annotated binding motif in the HOMER or JASPAR (2020 core vertebrate) motif databases, and TFs lowly expressed (mean FPKM < 0.8 across four replicates) in 2i-cultured wild-type mouse embryonic stem cells, based on RNA-seq data (GSE159937). This yielded 241 candidate target TFs.

Candidate spacer sequences for each target were drawn from a previously published, genome-scale mouse CRISPRa spacer library^32^. All library entries matching each target gene symbol were retrieved; for the small number of targets with no matching entry, the correct gene symbol for that TF in the spacer library was identified by cross-referencing gene symbol synonyms (Mouse Genome Informatics Homology database) and the retrieval repeated (e.g., *Trp63* in place of *Tp63*). One candidate (*Hoxd4*) had no corresponding entries in the spacer library and was excluded, leaving 240 targets with spacers available for selection.

For each of these 240 TFs, three top-ranked spacers were selected in total, with the specific spacers chosen according to the target’s annotated transcription start site (TSS) structure: for targets with separately annotated primary (P1) and secondary (P2) TSSs, the two top-ranked spacers targeting P1 and the top-ranked spacer targeting P2 were selected; for targets with spacers annotated as jointly targeting both TSSs (P1P2), the three top-ranked such spacers were selected; and for targets with only a single annotated TSS, the three top-ranked spacers against that TSS were selected. This yielded a total of 720 spacers across the 240 targets. Selected protospacer sequences were flanked with vector-specific cloning adapters (5′-atcttgtggaaaggacgaaacacc-3′ and 5′-gtttaagagctatgctggaaacagcatagcaagt-3′) and synthesized as single-stranded oligonucleotides (Integrated DNA Technologies) for cloning into the sgRNA expression vector.

#### Cloning of gRNA libraries into CROP-seq vector

Spacer sequences used for both the 240 TF RNA screen as well as the 60 TF co-assay screen were ordered from IDT as oPool Oligo Pools. Oligos were resuspended to 100µM and then serially diluted to 1 ng/uL, which was used as input for qPCR to append homology to an optimized CROP-seq backbone (Addgene #106280)^33,34^: PCR amplification was performed for 10 cycles using Kapa Biosystems HiFi Hotstart ReadyMix (KHF, Roche cat#KK2602) with forward primer “AH13 ATCTTGTGGAAAGGACGAAACA”, reverse primer “SD56 CTGTTTCCAGCATAGCTCTTAAAC” , a 63.5°C annealing temperature a 15 second extension, and purified using Zymo Research DNA Clean & Concentrator-5 kit. The CROP-seq vector was linearized with BsmBI-v2 (NEB) and agarose gel purified using a Zymoclean Gel DNA Recovery Kit (Zymo Research). Purified vector and insert were assembled via Gibson assembly using NEBuilder HiFi DNA Assembly Cloning Kit (NEB) and a ∼3:1 molar ratio of insert to vector. Assembled libraries were transformed into Stable Competent E. coli (NEB C3040H) and grown in liquid culture at 30°C; a small portion of transformants were also plated so that the number of colony forming units could be estimated to ensure a minimum 1,000X coverage of the library was achieved. Plasmid DNA was isolated using ZymoPURE Plasmid Miniprep Kit (Zymo Research) and subsequently packaged into lentivirus.

#### Lentivirus generation

Cloned CROP-seq gRNA libraries were packaged in lentivirus using the third generation packaging system following a protocol: The day before transfection, 2 million HEK293T cells were seeded on 5 10-cm dishes. For each transfection plate, 10µg of CROP-seq gRNA library was mixed with 5µg of pMDLg/pRRE^79^ (Addgene #12251), 2.5µg of pRSV-Rev^79^ (Addgene #12253), and 2.5µg of pMD2.G (Addgene #12259) in 600µL of DMEM, after which 50µL of FuGENE 6 transfection reagent (Promega cat#E2693) was added to the diluted plasmids and incubated for 20 minutes at room temperature before being added dropwise to each plate of cells. Approximately 4 hours after transfection, the media was replaced with fresh DMEM; 48 hours post transfection, lentivirus-containing media was collected from cells and collected again 60 hours post transfection (for a total of two collections). Viral media was concentrated using 5X PEG-it virus precipitation solution (System Biosciences cat#LV825A-1) following manufacturer instructions. For cDNA vectors, OmicsLink™ Expression-Ready ORF cDNA Vectors were used from GeneCopoeia in lentiviral pEZ-Lv236 vectors. These place the cDNAs of Gata6, Neurog2 and Fosl1 under the control of a CAG promoter and also include a puromycin resistance gene. Otherwise similar steps were followed as for the gRNA constructs.

#### Lentiviral transduction and multiplicity of infection estimations

To determine the volume of lentivirus to transduce cells that would result in a low multiplicity of infection, 250,000 mESC cells were seeded in every well of 2 gelatinized 6-well plates. The following day, medium was replaced with fresh 2i medium containing 0.1% protamine sulfate (8 mg/ml, Sigma P-45605), and each respective well was transduced with varying volumes of PEG-it concentrated lentivirus (ranging from 2.5µL to 50µL of virus per well). Approximately 8 hours after the initial transduction, each well was infected a second time using the same volume of lentivirus and protamine sulfate that had been applied in the first round of infection. The following day, each well of transduced cells was expanded into 2-wells of a gelatinized 6-well plate. Approximately 8 hours after expansion, media was replaced with fresh 2i media containing 2µg/mL puromycin and cells were selected for 48 hours after which point the surviving cells were counted for each lentivirus infection condition. To achieve a target MOI of <1, the lentivirus volume that yielded approximately ≤10% survival of infected cells after selection (relative to unselected controls) was chosen. These same infection conditions were then applied to cells used for single-cell experiments.

#### qRT-PCR to estimate target TF activation levels

For quantification of activation levels via qRT-PCR, cells were transfected or transduced with gRNAs expressed from the sgOpti backbone. RNA was isolated every 24h for up to 120h using the RNeasy Mini kit (Qiagen #74104) and converted into cDNA using SuperScript IV (ThermoFisher #18090010) and random hexamer primers. Primers for qPCR of target genes were designed to be situated within exons and have matching amplicon sizes and their efficiency was determined on a dilution curve of genomic DNA. To assess activation levels, target gene expression was compared to samples without guide RNA transfection/transduction and normalized to *Gapdh*.

#### Bulk ATAC-seq

We generated bulk ATAC-seq datasets following the Omni-ATAC protocol^80^. Briefly, isolated nuclei pellets were resuspended in freezing buffer^81^, flash frozen in liquid nitrogen and stored in -80°C. On the day of the experiment, nuclei were thawed in a 37°C water bath, pelleted at 500 x g for 5 min at 4°C, and repermeabilized in Omni lysis buffer (10 mM Tris-HCl pH 7.4, 10 mM NaCl, 3 mM MgCl2, 0.1% NP40, 0.1% Tween20, 0.01% Digitonin) on ice for 3 minutes, then neutralized with RSB-Tween (10 mM Tris-HCl pH 7.4, 10 mM NaCl, 3 mM MgCl2, 0.1% Tween20). 50,000 nuclei were used per tagmentation reaction and incubated at 37°C for 30 minutes. Tagmented DNA were purified using Zymo DNA Clean and Concentrate kit (Zymo Research, D4013), eluted in EB buffer (10 mM Tris), and amplified using 2x NEBNext HF PCR Mastermix (NEB, M0541L) and indexed Illumina primers using SYBR green to monitor amplification and preserve library complexity. Amplified DNA were purified using Zymo DNA Clean and Concentrate kit (Zymo Research, D4013) and run on a D5000 screentape tapestation (Agilent, 5057-5588 screentape, 5067-5589 reagents) to assess concentration and visualize nucleosome banding. Libraries were normalized to 2 nM, pooled equimolarly and loaded onto a Illumina NextSeq 500 High 75 cycle kit at 1.5 pM loading concentration with paired end sequencing (read1: 40 cycles, read 2: 40 cycles, and index 1: 10 cycles).

#### Cell and nuclei preparation and processing for 10x Genomics experiments

Cells were lentivirally transduced with respective gRNA libraries and puromycin selected as described above (see “Lentiviral transduction”). Three days after initiation of puromycin selection (*i.e.* four days after transduction), dead cells were removed by washing cultures three times with DPBS, and cells were harvested by incubation with Accutase for approximately 2-3 minutes, quenched with culture media, and pooled into a single conical tube before processing.

For the 240 TF 3’ scRNA-seq screen, whole cells were used as input on 12 lanes of 10x Genomic’s Chromium Single Cell 3ʹ v3 kit (target recovery of 10,000 cells per lane) following the respective manufacturers protocol.

For Multiome experiments, approximately 1 million cells were aliquoted for nuclei isolation following the 10x Genomics demonstrated protocol for nuclei isolation for Single Cell Multiome assays (CG000365, Rev. B). Nuclei were processed using 10x Genomic’s Chromium Next GEM Single Cell Multiome Kit following manufacturer’s protocol using 2 lanes and 16 lanes of the kit for the 10-TF pilot screen and 60-TF full screen, respectively.

#### gRNA enrichment from cDNA libraries via nested qPCR

For all single cell screens using 10x Genomics kits, gRNAs were captured via qPCR with 3 rounds of nested PCRs starting from purified amplified cDNA library. All PCRs were performed using Kapa Biosystems HiFi Hotstart ReadyMix (KHF, Roche cat#KK2602) in 50 µL reactions containing 1.5 µL of 10 µM forward and reverse primers and SYBR Green (Thermo Fisher #A25742) to monitor reactions in real time and prevent overamplification/saturation. Cycling conditions consisted of an initial denaturation at 95°C for 3 minutes, followed by repeated cycles of 98°C for 20 seconds, 65°C/65°C/72°C for 15 seconds, and 72°C for 50 seconds, with a final extension at 72°C for 1 minute. Reactions were purified using 1X Agencourt AMPure XP beads. Primers for the respective rounds of PCRs were:

*First PCR*: U6_Outer (5’-TTTCCCATGATTCCTTCATATTTGC-3’) + R1_PCR1 (5′-ACACTCTTTCCCTACACGACG-3′)*Second PCR*: U6_inner

(5′-GTCTCGTGGGCTCGGAGATGTGTATAAGAGACAGcTTGTGGAAAGGACGAAACAC-3′) + R1-P5 (5′-AATGATACGGCGACCACCGAGATCTACACTCTTTCCCTACACGACG-3′)

*Third indexing PCR*: Nextera P7 indexing primer

(5′-CAAGCAGAAGACGGCATACGAGAT(index)GTCTCGTGGGCTCGG-3′) + R1-P5 (5′-AATGATACGGCGACCACCGAGATCTACACTCTTTCCCTACACGACG-3′)

#### Sequencing run information

| Experiment | Library/sample | Instrument | Reagent kit | Run structure |
|---|---|---|---|---|
| Pilot Bulk ATAC-seq | *Fosl1*, *Gata6*, *Neurog2* activated (gRNA+cDNA) and WT bulk ATAC-seq | NextSeq 500 | High Output 75-cycle | 40/10/0/40 |
| Pilot Multiome | Transcriptome + gRNA association | NextSeq 500 | High Output 150-cycle | 28/10/10/90 |
| Pilot Multiome | ATAC-seq | NextSeq 2000 | P2 100-cycle | 50/8/24/49 |
| 240-TF scRNA-seq | Transcriptome + gRNA association | NextSeq 2000 | P3 100-cycle | 28/10/10/90 |
| 240-TF scRNA-seq | Transcriptome + gRNA association | NextSeq 2000 | P3 100-cycle | 28/10/10/90 |
| 60-TF Multiome | ATAC-seq | NextSeq 2000 | P3 100-cycle | 50/8/24/49 |
| 60-TF Multiome | ATAC-seq | NovaSeq 6000 | S2 100-cycle v1.5 | 50/8/24/49 |
| 60-TF Multiome | Transcriptome + gRNA association | NextSeq 2000 | P3 100-cycle | 28/10/10/90 |
| 60-TF Multiome | Transcriptome + gRNA association | NextSeq 2000 | P3 100-cycle | 28/10/10/90 |
| 60-TF Multiome | gRNA association | NextSeq 2000 | P2 100-cycle | 28/10/10/90 |

### Computational methods

Pilot bulk ATAC-seq data processing and analysis

Pilot bulk ATAC-seq datasets were generated for *Gata6*, *Neurog2*, and *Fosl1* activation conditions together with WT control samples. For each TF, both CRISPRa gRNA-mediated activation and cDNA overexpression conditions were profiled. For this proof-of-concept experiment, CRISPRa gRNA activation conditions and WT controls were profiled in biological duplicate, while cDNA overexpression conditions were profiled with a single replicate. Bulk ATAC paired end sequencing reads were demultiplexed with bcl2fastq (v2.20) and resulting FASTQ files were aligned to the mm10 mouse reference genome using BWA-MEM^82^ (v0.7.17) with the -M option and otherwise default parameters. Alignments were sorted and indexed using SAMtools^83^ (v1.19). Peaks were called independently for each bulk ATAC-seq sample using MACS2^84^ (v2.2.7.1) with paired-end BAM input (-f BAMPE), the mouse effective genome size (-g mm), model building disabled (--nomodel), ATAC-seq centering parameters (--shift -75 --extsize 150), summit calling enabled (--call-summits), and a p-value threshold of 0.01.

For each TF comparison, sample-specific MACS2 narrowPeak files from TF-activated and WT control bulk ATAC-seq samples were imported using rtracklayer (v1.62.0) and merged and reduced using GenomicRanges (v1.54.1) to generate a unified peak set. For GATA6 and NEUROG2, peak sets called from the corresponding pilot Multiome pseudobulked scATAC-seq profiles were also included in the unified peak set to enable direct comparison between bulk and single-cell-derived ATAC profiles (see “Data processing and analysis of pilot 10-TF Multiome dataset” below). Peaks were restricted to standard chromosomes (chr1–19, chrX, chrY, and chrM) using the keepStandardChromosomes function from GenomeInfoDb (v1.38.8), and regions overlapping the mm10 blacklist were removed. Bulk ATAC-seq BAM files were organized into QuasR^85^ (v1.42.1) sample files and loaded using qAlign with the mm10 genome and paired-end fragment orientation specified as paired = "fr". Fragment counts over the resulting consensus peak set were then quantified using qCount with strand-agnostic counting (orientation = "any").

Raw peak counts were normalized to counts per million mapped fragments for each sample. Fold-changes were calculated as log_2_-normalized CPM differences with a pseudocount of 1, averaging replicates where available. Differentially accessible peaks were defined as peaks with at least three-fold increased or decreased accessibility relative to WT controls. Peak-level concordance between samples was assessed using Pearson and Spearman correlations of normalized accessibility counts. Motif enrichment analyses on differentially accessible peaks were subsequently performed using monaLisa as described below (see “Motif enrichment in differentially accessible sites with monaLisa”).

#### Data processing and analysis of pilot 10-TF Multiome dataset

Pilot 10-TF Multiome libraries were processed using Cell Ranger ARC v2.0.0. Each replicate was processed independently with cellranger-arc count using the 10x Genomics mm10 Cell Ranger ARC reference (refdata-cellranger-arc-mm10-2020-A) and default parameters. Replicates were then aggregated using cellranger-arc aggr with depth normalization enabled. gRNA association libraries were processed separately using Cell Ranger v5.0.1 with ARC-v1 chemistry, the mm10 gene expression reference, and a feature reference containing expected gRNA sequences.

gRNA assignments were generated from the gRNA association BAM files using a custom Python script ^34^. Briefly, unmapped reads containing the sequence immediately upstream of the gRNA barcode were identified, and the downstream barcode sequence was extracted. Barcodes were corrected to a whitelist when possible, and gRNA–cell barcode associations were summarized for each cell barcode. Because gRNA recovery was relatively sparse in this proof-of-concept pilot dataset, associations were filtered permissively by removing associations supported by fewer than 100 reads without imposing an additional UMI-count cutoff, since downstream analyses primarily relied on aggregated signals expected to be robust to low levels of assignment noise.

Aggregated Multiome data were loaded into Signac^86^ (v1.13.0) using the filtered feature-barcode matrix and ATAC fragment file output by Cell Ranger ARC. Gene annotations were obtained from EnsDb.Mmusculus.v79 (v2.99.0) and converted to UCSC/mm10 chromosome naming. A Signac object was initialized from the gene expression matrix, and an ATAC chromatin assay was added using the peak matrix, fragment file, and gene annotations. gRNA assignments, replicate identity, and Scrublet doublet scores were added as metadata. Cells were filtered to retain high-quality singlets with mitochondrial RNA percentage below 20%, at least 1,000 RNA UMIs, 1,000–100,000 ATAC counts, and Scrublet doublet score below 0.2.

Using the filtered Signac object, peaks were called on subsets of TF-assigned cells and NTC cells using MACS2 (v2.2.7.1) as implemented through the Signac (v1.13.0) CallPeaks function. To generate pseudo-bulk ATAC BAM files, we generated cell barcode sets corresponding to each TF-assigned group and the NTC group from the filtered Signac object. For each Cell Ranger ARC replicate, the position-sorted ATAC BAM file was subsetted using samtools view with the CB cell barcode tag to retain reads derived from the corresponding selected cell barcode set. Resulting BAM files were indexed using samtools index.

These pseudobulked ATAC BAMs and peak sets were then included alongside bulk ATAC-seq BAMs for downstream consensus peak generation, fragment counting with QuasR^85^ (v1.42.1), normalization, and concordance analyses as described for the bulk ATAC-seq analysis described above.

#### Data processing for the 240-TF scRNA-seq dataset

Raw sequencing data were demultiplexed using cellranger (cellranger v5.0.1) mkfastq for transcriptome libraries and bcl2fastq (bcl2fastq v2.20) for gRNA association libraries. Transcriptome reads were aligned and quantified using cellranger (cellranger v5.0.1) count with default parameters, the mm10 Cell Ranger reference (refdata-gex-mm10-2020-A) and an expected cell recovery of 9,000 cells per sample. gRNA association libraries were processed separately using cellranger count (cellranger v2.1.1) with 10x Genomics 3′ v2 chemistry.

gRNA assignments were generated from the gRNA association BAM files using a custom Python script. Briefly, unmapped reads containing the sequence immediately upstream of the gRNA barcode were identified, and the downstream barcode sequence was extracted. Barcodes were corrected to a whitelist when possible, and gRNA-barcode associations were summarized for each cell barcode. Associations supported by fewer than 100 reads or fewer than 10 UMIs, or containing unprocessed barcode sequences, were removed.

Predicted doublet scores were calculated independently for each transcriptome library using Scrublet^87^ (v0.2.3) through Scanpy^88^ (v1.8.2). Transcriptome count matrices were then loaded into Seurat ^89^ (v5.1.0), merged across all 12 lanes, and annotated with sample lane identity, gRNA assignments, and Scrublet doublet scores. Cells with Scrublet doublet scores greater than 0.2 were removed. Cell-cycle scores were calculated in Seurat using mouse orthologs of canonical human S-phase and G2/M marker genes, and mitochondrial read percentages were calculated using genes matching the ^mt- prefix.

The filtered Seurat object was converted to a Monocle3^90^ (v1.3.5) cell data set for further processing and dimensionality reduction. Cells with greater than 10% mitochondrial reads or fewer than 1,000 UMIs were excluded. The dataset was preprocessed using 100 principal components, and batch effects were corrected across replicates using align_cds. UMI count, S-phase score, and G2/M score were included as residual covariates during alignment to regress out cell-cycle phase and UMI counts. Dimensionality reduction was then performed with UMAP using the aligned representation, with 30 nearest neighbors, a minimum distance of 0.1, and three output dimensions. Cells were clustered using Monocle3 with default cluster_cells parameters, and the resulting processed cell data set was saved for downstream analyses.

#### Selection of top 60 TFs to follow up on in secondary Multiome screen

Candidate TF perturbations were prioritized from the 240-TF scRNA-seq screen by testing whether cells assigned a given gRNA were non-randomly distributed in low dimensional UMAP space, i.e. sharing transcriptional cell states. Briefly, a sparse binary matrix was constructed indicating the presence or absence of each gRNA assignment in each cell, and a Monocle3 cell data set object was generated using the previously computed UMAP coordinates from the Monocle3 processed scRNA-seq data. Moran’s I spatial autocorrelation statistics were then calculated for each gRNA using Monocle3 graph_test on a k-nearest-neighbor graph (neighbor_graph = "knn") with k = 500. This analysis identified gRNAs whose assigned cells formed localized enrichments in transcriptional state space relative to the background cell population. The top 60 TFs were selected based on the most significant gRNA-level Moran’s I q-values. As an orthogonal approach that did not rely on the low-dimensional UMAP representation, TFs were also ranked using lochNESS neighborhood enrichment scores^39^ calculated from the RNA nearest-neighbor graph in PCA space (see “*lochNESS filtering*” below). For each TF, the mean lochNESS score across assigned cells was multiplied by the number of assigned cells to account for the dependence of raw lochNESS scores on gRNA frequency, and the top 60 TFs were selected based on this weighted score. Of the 60 TFs selected by Moran’s I, 47 were also among the top 60 ranked by this lochNESS scoring approach.

#### Data processing for the 60-TF 10x Genomics Multiome dataset

Raw sequencing data were demultiplexed using cellranger-arc (v2.0.2) mkfastq for Multiome libraries and bcl2fastq (v2.20) for gRNA-association libraries. Multiome libraries were processed using cellranger-arc count with the mm10 Cell Ranger ARC reference (refdata-cellranger-arc-mm10-2020-A). gRNA association libraries were processed separately using cellranger (v2.1.1) count with 10x Genomics 3′ v2 chemistry to generate BAM files for downstream barcode extraction. Samples were subsequently aggregated across replicates using cellranger-arc aggr with depth normalization enabled. gRNA assignments were generated as described for the previous screen (see “Data processing for the 240 TF scRNA-seq dataset” above) using a UMI and read count threshold of ≥4 UMIs and ≥150 reads per association.

Aggregated Multiome data were loaded into Seurat/Signac using the filtered gene expression matrix and ATAC fragment file produced by Cell Ranger ARC. Cells were filtered to retain those with 800–50,000 ATAC counts, 800–20,000 RNA UMIs, and less than 30% mitochondrial reads. gRNA and target assignments were added to cell metadata by matching corrected cell barcodes from the gRNA-association libraries to Multiome cell barcodes. Cell-cycle phase assignments were calculated using Seurat as described for the 240 TF RNA screen.

Because accessibility changes induced by individual TF perturbations may be present in only a subset of cells and therefore missed by peak calling on the full pooled dataset, peaks were called separately for each TF-assigned cell population. MACS2, as implemented through Signac’s CallPeaks function, was used to call peaks independently on cells assigned to each of the 60 TF perturbations, NTC cells, and cells with no detectable gRNA assignment. Peak sets from all TF-specific and control peak calls were concatenated into a single GRanges object, and overlapping genomic intervals were collapsed into a unified peak set using reduce from the GenomicRanges^91^ (v1.54.1) package. Peaks on nonstandard chromosomes and ENCODE blacklist regions were removed prior to downstream analyses.

A new binary cell-by-peak matrix was then generated from the aggregated unified peak set using Signac FeatureMatrix, thereby rescoring each cell for accessibility across the final merged set of peaks. This matrix was converted into a Monocle3 cell data set and processed using latent semantic indexing (LSI) dimensionality reduction. UMAP embeddings and clusters were generated from the LSI representation and transferred back into the Signac object as dimensional reduction slots. Neighbor graphs were constructed in Seurat using the Monocle3-derived LSI embeddings.

For RNA analyses, raw RNA counts were separately converted into a Monocle3 cell data set. Principal component analysis was performed using preprocess_cds with 100 dimensions, and RNA UMI counts were regressed using align_cds. UMAP embeddings and clusters were generated in Monocle3 and transferred back into the Seurat object. RNA data were subsequently normalized, variable features identified, and scaled in Seurat prior to neighborhood graph construction and clustering using the Monocle3-derived PCA embeddings. Finally, the aggregated TF-specific ATAC assay was TF-IDF normalized and subjected to singular value decomposition (SVD) using Signac for downstream chromatin accessibility analyses.

#### lochNESS filtering

To identify high-confidence gRNA(+) cells for downstream analyses, we implemented a neighborhood enrichment–based filtering strategy adapted from lochNESS^39^. For each TF, cells assigned a corresponding gRNA (including cells assigned multiple gRNAs) were labeled as “KO”, while all remaining cells were labeled as “WT”. When lochNESS was run on the NTC population, cells assigned an NTC gRNA (but no targeting gRNAs) were instead classified as “KO”, with all remaining cells treated as “WT”.

Neighborhood enrichment scores were calculated using the weighted ATAC nearest-neighbor graph derived from low-dimensional chromatin accessibility embeddings. For each cell, edge weights connecting the cell to neighboring KO cells were summed and scaled by the neighborhood size parameter k. This local KO-neighbor enrichment was then normalized by the global frequency of KO cells in the dataset, with 1 subtracted so that a score of 0 reflects the enrichment expected under random cell mixing. Scores greater than 1 therefore indicate local KO enrichment more than two-fold above expectation. The neighborhood size parameter was defined as k = round(0.5 × √N), where N is the total number of cells in the dataset. Cells assigned to a TF perturbation with a neighborhood enrichment score greater than 1 were retained as high-confidence gRNA(+) cells for downstream analyses.

#### Differential expression testing

Unless otherwise noted, for differential expression (and accessibility) testing, cells assigned multiple TF-targeting gRNAs were included in the TF cell population for each corresponding TF’s analysis, whereas the NTC population was restricted to cells that were only assigned NTC gRNAs and no TF-targeting gRNAs. We confirmed overall UMI counts are similar between these populations.

#### Standard differential expression testing

Differential expression testing was performed using the Wilcoxon rank-sum test implemented in Seurat’s FindMarkers function. Unless otherwise noted, genes were required to be detected in ≥5% of cells in at least one of the populations being compared. P-values were adjusted for multiple testing using the Benjamini–Hochberg method. Additional fold-change thresholds were applied depending on the analysis and are indicated in figure legends where relevant. For analyses quantifying the number of differentially expressed genes, we generally required an absolute average log_2_ fold-change ≥1 in addition to statistical significance and the detection-frequency threshold unless otherwise noted.

#### UMI-matched differential expression testing

For the UMI-matched differential expression analysis shown in **Supplementary Fig. 12**, we controlled for differences in total RNA UMI counts between TF-targeting and NTC cells. For each TF independently, cells were binned according to total RNA UMI count using 30 bins spanning the combined range of UMI counts across the TF-targeting and NTC populations. NTC cells were then sampled within each bin in proportion to the number of TF-targeting cells in that bin, yielding NTC populations approximating the total cell numbers and UMI-count distributions of the corresponding TF-targeting populations. Differential expression between TF-targeting cells and the resulting UMI-matched NTC population was performed as described above. For visualization, genes were required to have a Benjamini–Hochberg-adjusted p < 0.05, ≥1.5-fold average increase in expression, and expression detected in ≥5% of TF-targeting cells.

#### Nearest-neighbor differential expression testing

For each of the 60 TFs, target cells were defined using one of two schemes: (1) all cells assigned a single gRNA targeting that TF ("gRNA" subset), or (2) the subset of gRNA-assigned cells further filtered by lochNESS score, retaining only cells with a raw mutant score greater than 1 ("lochNESS gRNA" subset).

For each target cell, we identified a matched background cell by querying its 50 nearest neighbors in a shared PCA embedding (RNA- or ATAC-derived PCA, specified per run) using a k-d tree search (FNN v1.1.4; get.knnx). Neighbors already assigned to the target set, or already selected as a background match for a previous target cell, were excluded, and the nearest remaining eligible neighbor was assigned as that cell’s paired background control.

With target and matched-background groups assigned, we performed DE testing on RNA-assay normalized expression using Seurat’s “FindMarkers” with a Wilcoxon rank-sum test, disabling default minimum percent-expressed and log-fold-change pre-filters (min.pct = -Inf, logfc.threshold = -Inf).

For each TF, we generated a null distribution by repeating the identical NN-matched DE procedure using randomly sampled non-targeting-control (NTC) cells in place of true target cells. For each TF, we drew a number of NTC cells equal to the number of that TF’s target cells (and repeated this sampling 5 independent times) and computed matched-background NN-DE statistics identically to the target analysis, yielding 5 independent null p-value sets per TF.

Target gene-level p-values were recalibrated against the pooled null distribution (all 5 NTC-matched iterations combined) by computing an empirical p-value for each TF as the fraction of null p-values less than or equal to the observed target p-value. The empirical p-values were sorted, compared against the standard BH critical value line (fdr × rank/m), and the largest empirical p-value at or below its corresponding critical value was used to identify the largest passing rank. The corresponding nominal (i.e., unrecalibrated) target p-value at that rank was used as a per-TF significance cutoff, and applied to the full target gene list together with a minimum effect-size filter (average log_2_ fold-change > 0.1) to call significantly differentially expressed genes. TFs for which no gene passed the empirical FDR threshold were assigned no significant genes.

This procedure was performed independently for the "gRNA" and "lochNESS gRNA" target-cell populations.

### Differential accessibility testing with DESeq2

Differential accessibility analyses were performed using a bootstrap pseudobulk strategy implemented using DESeq2^92^ (v1.42.1). For each target TF, cells assigned to the corresponding gRNA and passing lochNESS-based filtering criteria were treated as the perturbed cell population, while cells assigned as NTC were used as background controls. To generate pseudobulk replicates, 100 bootstrap iterations were performed for each TF perturbation. In each iteration, 80% of perturbed cells were randomly sampled without replacement and aggregated by summing chromatin accessibility counts across peaks to generate a pseudobulk accessibility profile. In parallel, an equal number of NTC cells were randomly sampled and pseudobulked using the same procedure. This resulted in paired perturbed and control pseudobulk accessibility matrices for downstream differential accessibility testing.

Differential accessibility testing was performed using DESeq2, treating each bootstrap iteration as a replicate. Pseudobulk count matrices were processed using default DESeq2 settings with target versus NTC specified as the design variable. Because treating bootstrap iterations as independent replicates inflates statistical significance, empirical false discovery rate (FDR) thresholds were estimated using matched-control analyses based entirely on NTC cells. Briefly, for each TF, a number of NTC cells equal to the number of TF cells was randomly sampled from the overall NTC population and treated as a “pseudo-target” group, while the remaining NTC cells served as the outgroup. Bootstrap pseudobulking and DESeq2 testing were then performed identically to the primary analysis. This procedure was repeated across five independent random seeds to generate empirical null distributions of differential accessibility statistics.

Empirical FDR thresholds were determined separately for each TF perturbation. Peaks were first filtered independently within the TF and null-control analyses to retain only those with an absolute log_2_ fold-change greater than 1. For a given adjusted p-value threshold t, an empirical FDR was calculated as the number of null-control peaks passing threshold t divided by the number of observed TF peaks passing the same threshold. The largest adjusted p-value threshold corresponding to an empirical FDR ≤ 5% was identified for each null-control run, and the mean threshold across runs was used as the final empirical significance cutoff for that TF perturbation. Peaks with log_2_ fold-change ≥ 1 and adjusted p-values below this empirical threshold were classified as differentially gained (DG), while peaks with log_2_ fold-change ≤ -1 and adjusted p-values below threshold were classified as differentially lost (DL), with differentially accessible peaks comprising the union of both.

### Visualization of DG and DL sites in UMAP embedding

To visualize relationships between DG and DL called across TFs, the binarized single-cell ATAC count matrix was subsetted to all DG and DL sites called for all TFs and transposed. Dimensionality reduction was performed using LSI followed by UMAP with Monocle3 (v1.3.5) on the transposed matrix such that each observation represents a peak. To assign DA sites to peak modules (**Fig. 3E**), peaks were assigned to mutually exclusive groups in the following order of priority: (i) TF-specific groups (Cdx2, Sox7, Neurod1, Sox10, Pax3, and Sox5), comprising sites called as DG exclusively for the indicated TF (for Sox5, including sites DG for Sox5 and/or Nrf1 but no other TF; see **Supplementary Fig. 6G-H**); (ii) TFAP comprising sites DG for one or more trophoblast-associated TFs (Elf5, Grhl1, Gata1, Gata5, Gata6, Ovol1) and no other TFs; (iii) FOS/JUN, comprising remaining DG sites not assigned to the above categories; and (iv) pluripotency, comprising sites called as DL for any TF and not called as DG for any TF.

### Overlap with different peak sets (ChIP-seq, EB ATAC)

To compare differentially accessible peak classes with independent chromatin datasets, we quantified overlap with external ChIP-seq, DNase-seq, and ATAC-seq peak sets. Histone ChIP-seq peak sets for ES-E14 mESCs were obtained from ENCODE for H3K4me1 (ENCSR000CGN; ENCFF158GBZ), H3K4me3 (ENCSR000CGO; ENCFF993IIG), H3K9ac (ENCSR000CGP; ENCFF668UBL), H3K27ac (ENCSR000CGQ; ENCFF519QMV), H3K9me3 (ENCSR000ADM; ENCFF925BSH), and H3K36me3 (ENCSR000CGR; ENCFF362DZS)^42–44^.

ChIP-seq narrowPeak files were obtained for the indicated histone modifications, and genomic intervals were converted from BED coordinates to GRanges. For each accessibility peak, overlap with each biochemical peak set was assessed using findOverlaps from the Bioconductor GenomicRanges (v1.54.1) package. A peak was considered overlapping a given biochemical mark if it shared at least 1 bp with at least one peak from that dataset. When multiple biochemical peaks overlapped the same accessibility peak, the overlapping interval covering the largest fraction of the accessibility peak was retained for annotation.

For each accessibility peak class, we calculated the percentage of peaks overlapping at least one peak for each biochemical mark. We additionally calculated the percentage of peaks overlapping any of the assayed histone-mark peak sets. For comparison with bulk ATAC-seq, peaks from two control replicates were combined and overlapping intervals were merged using reduce from GenomicRanges (v1.54.1) before overlap analysis. DNase-seq peaks were included as an additional reference peak set. The resulting matrix of percentages was hierarchically clustered across both accessibility peak classes and biochemical marks using Euclidean distances and complete-linkage hierarchical clustering as implemented by pheatmap (v1.0.12), and displayed as a clustered heatmap.

For comparison with mouse embryoid body (mEB) chromatin accessibility data from Lalanne et al.^40^, pseudobulk accessibility profiles were obtained from GSE217683. Each mEB ATAC-seq peak was assigned to the cell type with the highest accessibility if accessibility was at least eight-fold higher (log₂ fold-change ≥3) than the mean across all other cell types; remaining peaks were classified as non-specific. Overlap of DG, DL, non-DA, and GC-matched background regions with each mEB peak category was determined using findOverlaps. Enrichment of DG, DL, and GC-matched regions relative to non-DA sites was assessed using one-tailed hypergeometric tests, with p-values adjusted using the Benjamini–Hochberg method.

### Guide RNA enrichment in Louvain clusters

To identify transcription factor perturbations enriched in specific ATAC Louvain clusters, we compared the representation of cells assigned a gRNA for each TF against cells assigned to non-targeting control (NTC) guides. For each Louvain cluster and each target, cells were divided into those within the cluster and those outside the cluster. A 2 × 2 contingency table was then constructed using the number of target-guide cells and NTC-guide cells observed inside versus outside the cluster. Guide enrichment was tested using a chi-square test. Enrichment effect size was quantified as the ratio between the fraction of target-guide cells assigned to the cluster and the fraction of NTC cells assigned to the same cluster. P-values were adjusted across all tested targets within each cluster using the Benjamini–Hochberg procedure.

### Motif enrichment in differentially accessible sites with monaLisa

Motif enrichment analyses were performed using monaLisa^93^ (v1.8.1) together with vertebrate position weight matrices (PWMs) obtained from JASPAR2020. Sites called as differentially accessible were converted to genomic ranges and resized to a fixed width of 500 bp centered on the original peak midpoint prior to sequence extraction. Genomic DNA sequences corresponding to these resized regions were extracted from the BSgenome.Mmusculus.UCSC.mm10 (v1.4.3) mm10 reference genome.

Motif enrichment testing was performed using the calcBinnedMotifEnrR function in monaLisa with genome-sampled GC-matched background regions (background = "genome"). Background sequences were sampled from the full mm10 genome with a genome oversampling parameter of 2. Motif enrichment analyses were run serially using a fixed random seed for reproducibility. monaLisa enrichment statistics including log_2_ enrichment (log_2_enr) and adjusted p-values were extracted from the resulting SummarizedExperiment object for visualization.

### Pseudotime trajectory of the ExEn-like cell state

To construct a pseudotime trajectory of the ExEn-like cell state, we jointly considered transcriptional and chromatin accessibility features that distinguished ExEn-like cells from non-targeting control (NTC) cells. ExEn-like cells were defined as cells belonging to Louvain cluster 20 based on the expression patterns of canonical ExEn marker genes (e.g., Gata4, Gata6, Lama1 and Dab2)^37^. Differentially expressed genes were identified between ExEn-like and NTC cells using a Wilcoxon rank-sum test implemented with FindMarkers in Seurat. Genes with an average fold-change ≥3-fold, expression detected in ≥5% of ExEn-like cells, and Benjamini–Hochberg-adjusted p < 0.01 were retained. Differentially accessible regions were similarly identified using FindMarkers applied to the chromatin accessibility data and were required to have an average log_2_ fold-change >2, Benjamini–Hochberg-adjusted p < 0.001, and accessibility detected in <10% of NTC cells. Genes corresponding to TFs whose targeting gRNAs were enriched in the ExEn-like population (*Gata6*, *Gata1*, *Sox7*, and *Hnf1b*) were excluded from the transcriptional feature set to prevent their direct CRISPRa-mediated induction from driving the trajectory. Chromatin accessibility features within ±1 kb of the transcription start sites of these TFs were likewise excluded, although no differentially accessible regions met this criterion.

For trajectory inference, log-normalized RNA expression values and TF-IDF-normalized chromatin accessibility values for the selected features were combined across ExEn-like and NTC cells. Features were scaled to zero mean and unit variance, followed by principal-component analysis using 10 components. A nearest-neighbor graph was constructed from these components using correlation distance, and diffusion components were calculated using Scanpy. The root of the trajectory was selected from NTC cells as the cell closest, by Euclidean distance in PCA space, to the centroid of all NTC cells. Diffusion pseudotime was then calculated using sc.tl.dpt.

For visualization of individual transcriptional and chromatin accessibility features along the resulting trajectory, cells were ordered by diffusion pseudotime and normalized feature values were smoothed using a centered rolling mean across 300 cells. Smoothed values for each feature were independently rescaled from 0 to 1.

### Co-motif enrichment analysis with chromVAR and epiwraps

#### Without stratification by TFAP2C expression

To assess enrichment of individual TF motifs and co-motif combinations across TF conditions, pseudobulk chromatin accessibility profiles were generated and analyzed using chromVAR-based deviation scoring. Analyses were performed using the epiwraps package ^94^ together with chromVAR^48^ (v1.24.0), motifmatchr (v1.24.0), and JASPAR motif annotations.

Pseudobulk groups were generated from lochNESS-filtered TF cells using the peakPbCountsSE function from epiwraps. Cells were grouped by perturbation identity and further split by their individual gRNAs, producing replicate-like pseudobulk accessibility profiles per guide for each TF (n = 3) and NTC population (n = 20). Analyses were performed using the same 60-TF ATAC peak set used in earlier analyses; peaks were resized to a fixed width of 400 bp centered on the original peak midpoint to facilitate chromVAR background matching. Pseudobulk accessibility counts were generated directly from the aggregated ATAC fragment file outputted by cellranger-arc aggr using the mm10 reference genome.

Motifs were obtained from the JASPAR2024^95^ CORE non-redundant vertebrate motif collection and filtered to retain only motifs corresponding to the curated motif set used in previous analyses based on JASPAR2020^96^ annotations. Motif occurrences within accessible regions were identified using motifmatchr::matchMotifs. chromVAR deviation scores were computed using computeDeviationsFaster from epiwraps, using GC content- and accessibility-matched background peak sets generated with chromVAR::getBackgroundPeaks (n iterations = 200) from the pseudobulk accessibility matrix. Background peak assignments were precomputed and reused across downstream deviation analyses. Differential motif accessibility between TF groups and NTC controls was assessed using linear modeling with limma^97^ (v3.58.1). For each TF perturbation, moderated t-statistics and Benjamini–Hochberg adjusted p-values were computed for all motifs.

To evaluate enrichment of motif co-occurrence patterns, motif pairs of interest (shown in **Supplementary Fig. 16**) were predefined, and co-occurring motif instances were identified using the motifCoOccurence function from epiwraps, requiring motif pairs to occur within 5–100 bp of one another (i.e. minDist=5L, maxDist=100L). chromVAR deviation scores were then recomputed for these co-motif feature sets using the same background model and pseudobulk accessibility matrix described above.

Differential accessibility testing for co-motif features was performed using the same limma framework applied to single motifs. The number of genomic regions contributing to each co-motif feature was retained and reported alongside statistical results to facilitate interpretation of co-motif enrichment estimates derived from small numbers of sites.

#### Motif co-occurrence enrichment binned by co-motif spacing

To test whether co-motif accessibility enrichment varied with spacing between the two motifs, we repeated the co-motif analysis using motifCoOccurence’s quantile-binning mode (nDistQuantiles = 20, maxDist = 1000 bp) on the same candidate motif pairs. Peaks were padded to capture motif instances up to 1000 bp beyond their original boundaries, and for each pair, peaks containing at least one instance of each motif were assigned the minimum center-to-center distance between the two. These distances were divided into 20 bins with approximately equal numbers of co-occurring instances, using pair-specific quantile breakpoints of the observed distance distribution rather than fixed-width intervals to keep the number of contributing sites (and thus the resulting chromVAR deviation ranges) comparable across bins.

chromVAR deviations were then recomputed for each of the 20 distance-binned co-motif features per pair, using the same background peak sets and pseudobulk accessibility matrix described above, merged with the single-motif deviation matrix, and tested for differential accessibility between each TF perturbation and NTC controls using the same limma framework, with the number of contributing sites retained per bin to aid interpretation.

#### Stratification by TFAP2C expression

To examine whether co-motif accessibility enrichment was modulated by *Tfap2c* expression, we repeated the chromVAR-based analysis after additionally stratifying pseudobulk groups by *Tfap2c* detection status. Cells were classified as TFAP2C+ or TFAP2C− based on presence or absence of detectable *Tfap2c* expression. Pseudobulk profiles were then generated for each combination of TF identity (or NTC), individual gRNA, and TFAP2C detection status, using the same peakPbCountsSE-based pipeline, 60-TF ATAC peak set, JASPAR-derived motif set, precomputed background peak sets, and pseudobulk fragment aggregation described above.

Single-motif chromVAR deviations were computed as before. For co-motif enrichment, the same predefined candidate motif pairs were tested for co-occurrence using motifCoOccurence, with co-occurring instances required to fall within 5–100 bp of one another (minDist=5L, maxDist=100L). Co-motif deviations were recomputed and merged with the single-motif deviation matrix as described above.

Differential accessibility was assessed using a linear model in limma in which cells were grouped by the combination of TFAP2C status (+/−) and perturbation status (CRISPRa target vs NTC), followed by moderated t-tests on a set of pre-specified contrasts: the overall effect of each perturbation averaged across TFAP2C+ and TFAP2C− pseudobulks ("CRISPRa"); the perturbation effect estimated separately within TFAP2C+ and TFAP2C− cells ("CRISPRa_in_pos" / "CRISPRa_in_neg"); the baseline TFAP2C+ vs TFAP2C− difference among NTC control cells ("TFAP2C_in_ctrl"); and the interaction contrast testing whether the perturbation’s effect differed between TFAP2C+ and TFAP2C− cells (i.e., a difference-of-differences test). Benjamini–Hochberg-adjusted p-values were computed for each contrast, for each motif and co-motif feature, and for each TF perturbation relative to NTC.

This analysis was performed using cells with a single gRNA assignment, and separately in a version that allowed the inclusion of cells assigned multiple gRNAs. In the latter case, each ambiguous cell was resolved to a single gRNA by comparing the individual cell counts of the specific candidate gRNAs detected in that cell and assigning the cell to whichever of those specific gRNAs had fewer cells.

## SUPPLEMENTARY FIGURES

**Figure S1.**
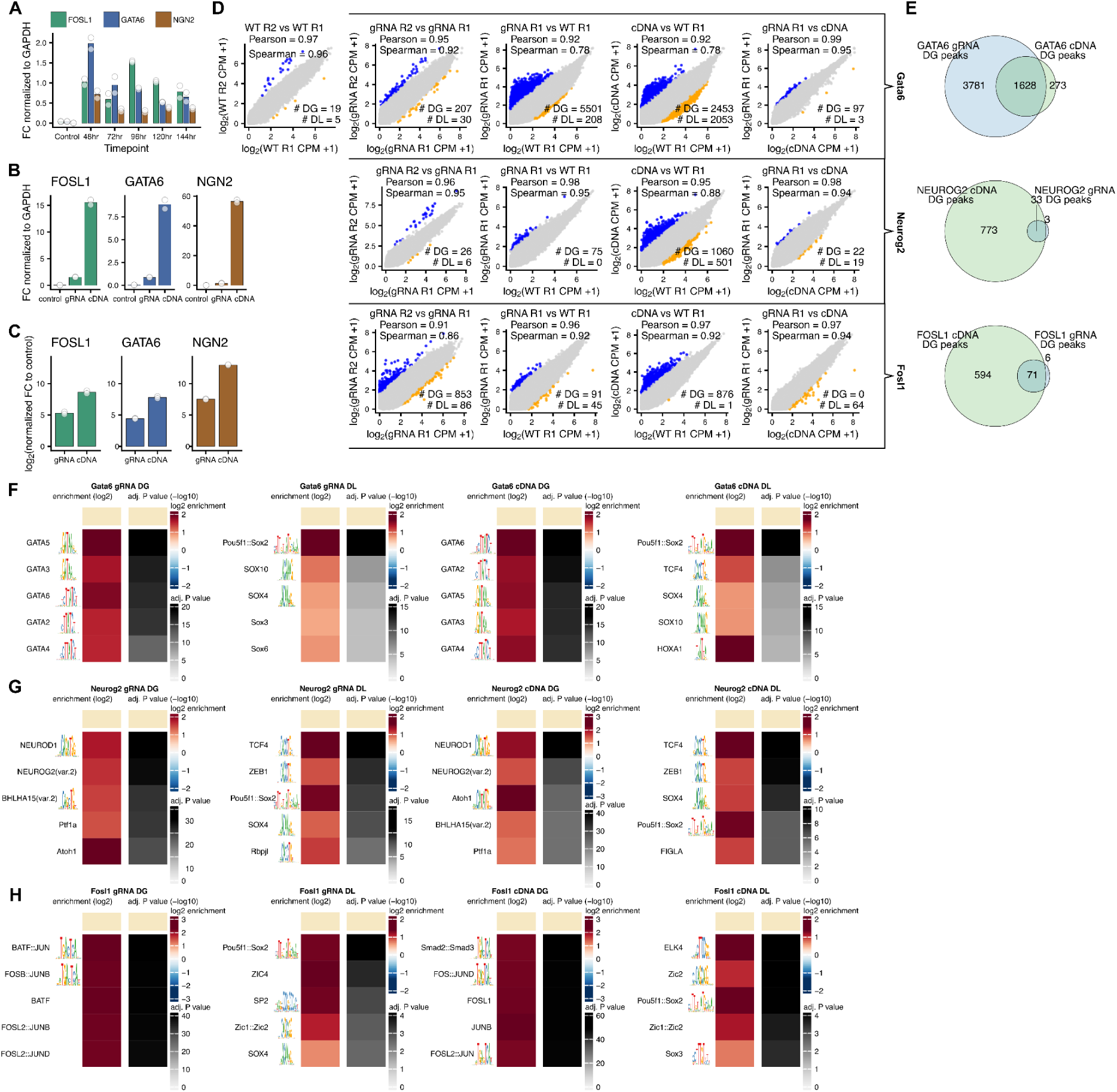
Comparison of TF activation and chromatin accessibility responses to CRISPRa and cDNA overexpression. **(A)** qPCR time series of CRISPRa activation at 48, 72, 96, 120 and 144 hrs after transduction of targeting guides or mock transduction. Values are normalized to *Gapdh* in the same sample, for each of two biological replicates. **(B)** qPCR activation levels for mock, gRNA and cDNA constructs relative to *Gapdh* in the same sample, for *Fosl1*, *Gata6* and *Neurog2* at 96 hrs post-transduction, for each of two biological replicates. **(C)** As in **B**, after normalizing to mock transductions. **(D)** Correlation plots comparing bulk chromatin accessibility (log_2_ CPM + 1) across replicates (R1, R2) and conditions (CRISPRa/gRNA or cDNA) for *Gata6* (top row), *Neurog2* (middle row) and *Fosl1* (bottom row) across merged peaks, each point a peak. Colors denote differentially accessible peaks. Because cDNA experiments lack replicates, DG and DL sites here were defined by fold-difference (blue: ≥3-fold increase; yellow: ≥3-fold decrease) of the first listed sample of each comparison relative to the second. Pearson and Spearman coefficients are shown for each comparison. Columns correspond, from left to right, to WT vs WT, gRNA vs gRNA, gRNA vs WT, cDNA vs WT, and gRNA vs cDNA. **(E)** Venn diagrams showing overlap between DG peaks in bulk gRNA and cDNA samples for *Gata6* (left), *Neurog2* (middle) and *Fosl1* (right). Peaks were called DG if exhibiting a ≥3-fold increase in normalized accessibility (CPM) relative to WT bulk ATAC controls. For gRNA samples, peaks were called in both replicates and merged. Only one replicate was available for cDNA samples. **(F-H)** monaLisa motif enrichment for *Gata6* **(F)**, *Neurog2* **(G)** and *Fosl1* **(H)**. For each TF, heatmaps show, from left to right, gRNA vs WT DG sites, gRNA vs WT DL sites, cDNA vs WT DG sites, and cDNA vs WT DL sites. DG and DL sites were the top 200 peaks ranked by fold-change relative to WT, without a significance or minimum fold-change threshold; sequences ±250 bp around peak midpoints were compared to a GC-matched genomic background. The top five enriched motifs by −log_10_ adjusted p-value are shown.

**Figure S2.**
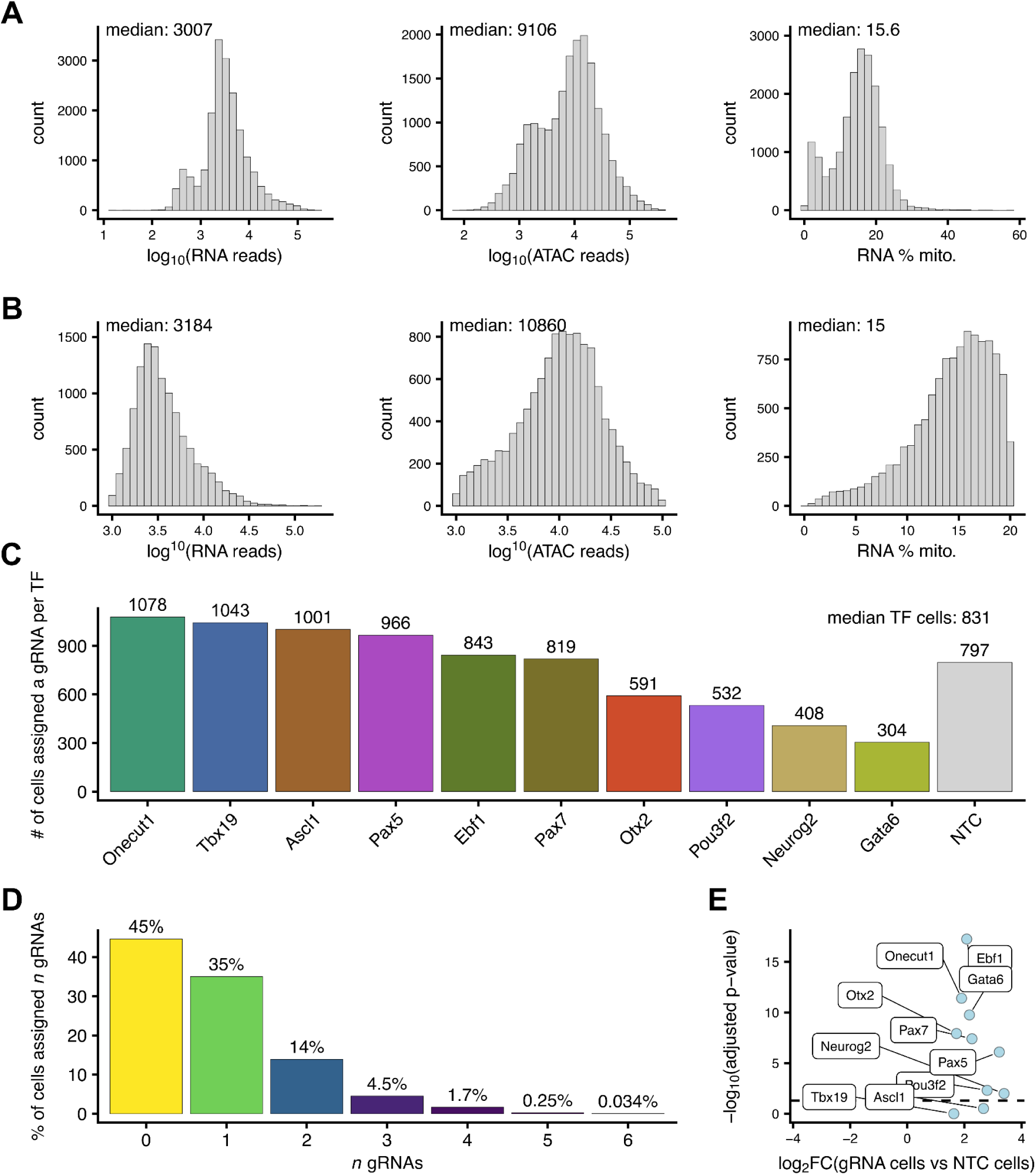
Pilot sc-multiome screen with pooled CRISPRa-activated TFs. (**A**) From left to right, distributions of per-cell RNA UMIs, unique ATAC fragments, and RNA mitochondrial read percentages, before filtering. (**B**) As in **A**, after filtering for cells with mitochondrial read percentage ≤20% and RNA and ATAC counts ≥1,000 (n = 11,615 cells). (**C**) Bar plots indicating the number of cells assigned a gRNA for each targeted TF (including cells with multiple gRNAs assigned) and the number assigned an NTC gRNA (single-gRNA assignments only). (**D**) Bar plots showing the percentage of cells assigned no gRNA, a single gRNA, or multiple gRNAs. (**E**) Fold change in expression of each target TF in cells bearing the corresponding gRNA versus NTC gRNA(+) cells.

**Figure S3.**
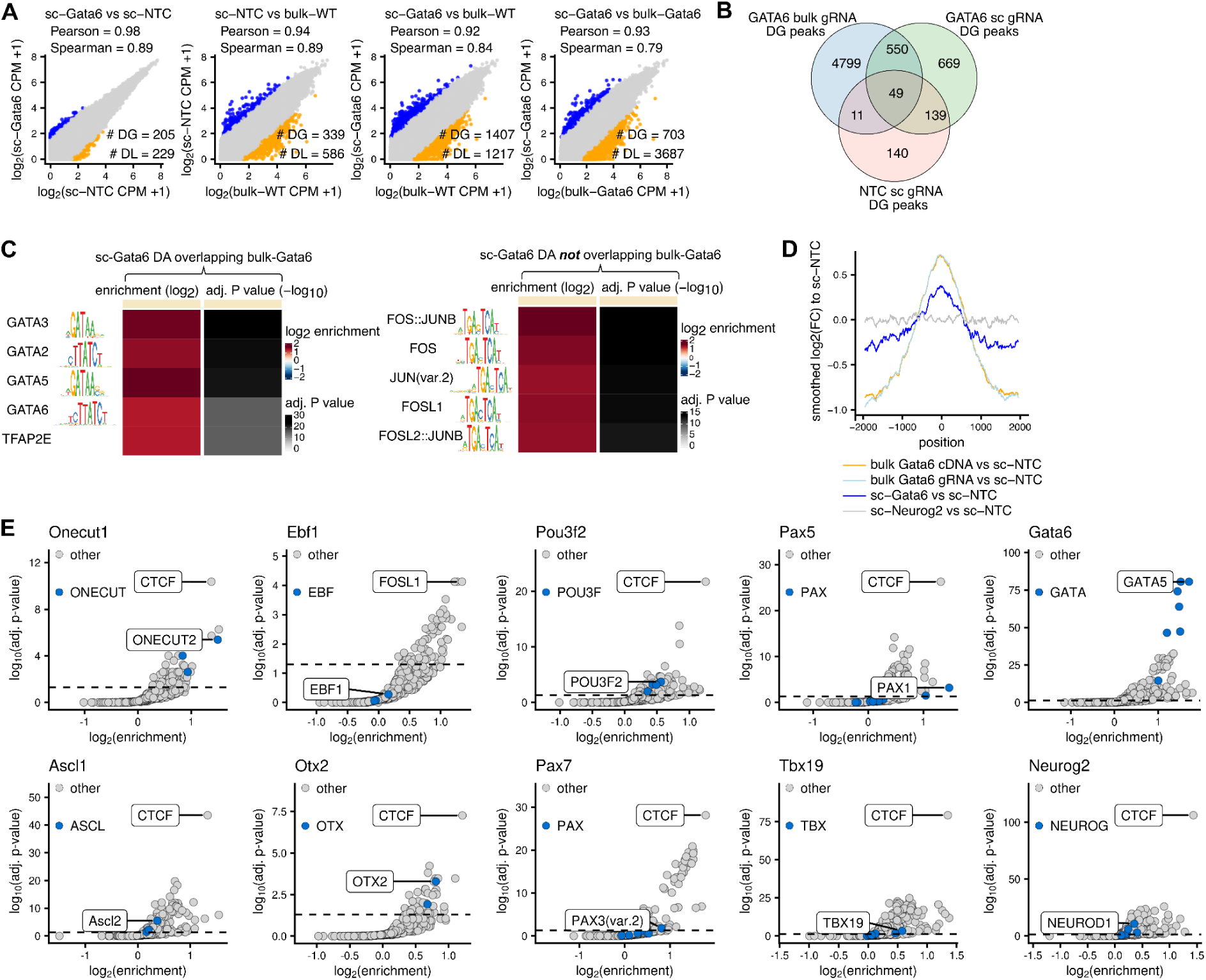
Shared signal between sc-multiome profiles and bulk ATAC-seq for CRISPRa-activated TFs. **(A)** Correlation plots comparing chromatin accessibility (log₂ CPM + 1) across pseudobulked *Gata6* gRNA(+) cells from the pilot sc-multiome screen, pseudobulked NTC gRNA(+) cells from the pilot sc-multiome screen, and bulk ATAC-seq of WT and *Gata6* CRISPRa-activated cells. Each point represents a single peak; colors denote peaks classified as differentially accessible. Because not all conditions have replicates, DG and DL sites here were defined by fold-difference (blue: DG ≥3-fold increase, yellow: DL ≥3-fold decrease) of the first listed sample of each comparison relative to the second. Pearson and Spearman correlation coefficients are shown for each comparison. **(B)** Venn diagram of overlap between the DG peaks in **A** for pseudobulked *Gata6* gRNA(+) sc-multiome cells, pseudobulked NTC gRNA(+) sc-multiome cells, and bulk *Gata6* gRNA ATAC samples. **(C)** monaLisa motif enrichment for DG peaks in pseudobulked *Gata6* gRNA(+) sc-multiome cells that overlapped bulk *Gata6* gRNA DG peaks (left) and for those that did not (right). Sequences ±250 bp around peak midpoints were compared to a GC-matched genomic background. The top five enriched motifs by −log₁₀ adjusted p-value are displayed. **(D)** Chromatin accessibility centered on DG sites called in bulk *Gata6* gRNA samples, showing smoothed fold change in normalized accessibility relative to single-cell NTC cells, for bulk *Gata6* cDNA, bulk *Gata6* gRNA, single-cell *Gata6* gRNA and single-cell *Neurog2* gRNA. Signals were averaged across DG sites, normalized per region and per million mapped reads, and smoothed by rolling mean.

**Figure S4.**
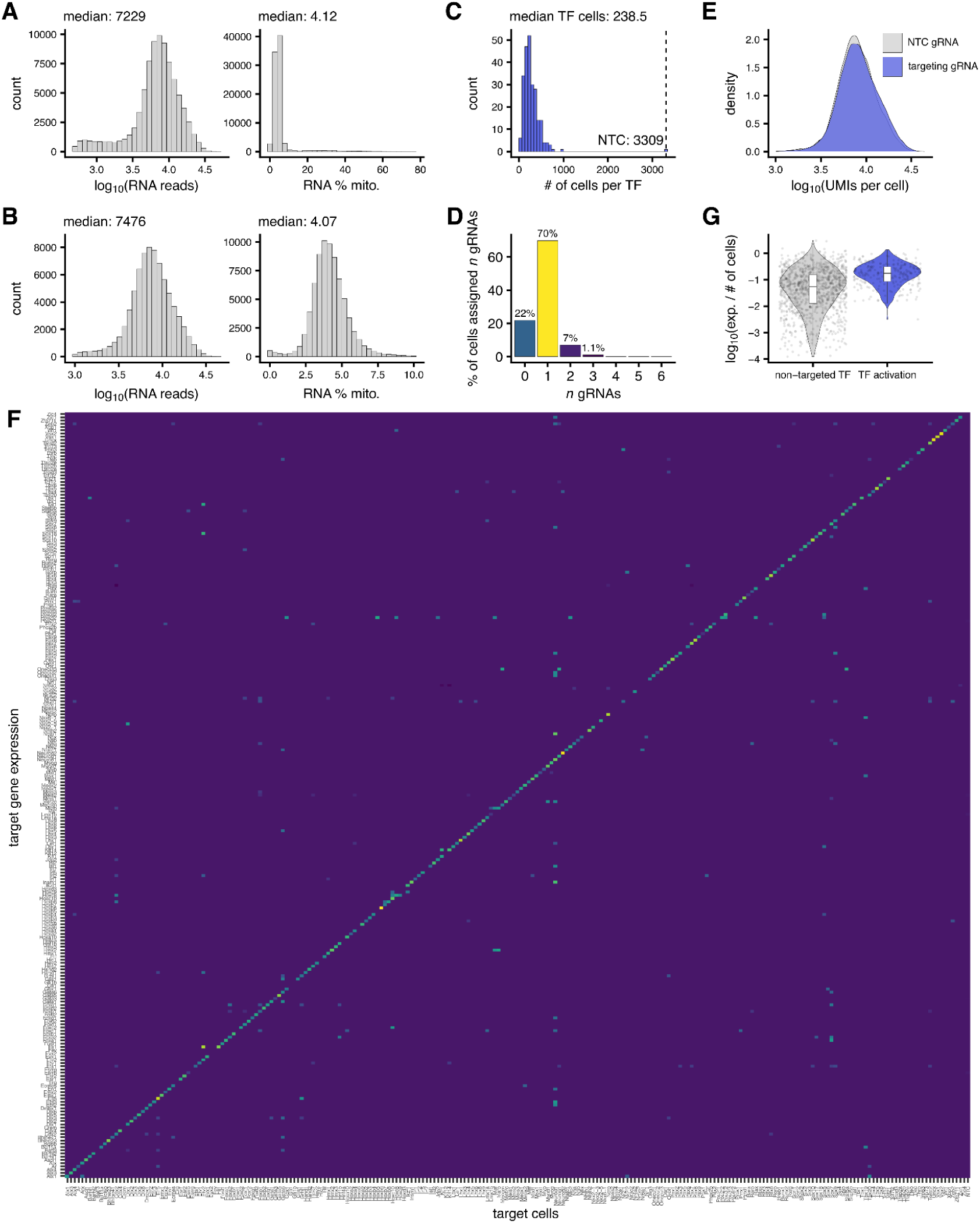
TF activation to physiological expression levels in the 240-TF scRNA-seq screen. **(A)** From left to right, distributions of per-cell RNA UMIs and RNA mitochondrial read percentages, before filtering. **(B)** As in **A**, after filtering for cells with mitochondrial read percentage ≤10% and RNA counts ≥1,000 (n = 75,340 cells). **(C)** Histogram showing the number of cells assigned a gRNA for each targeted TF (including cells with multiple gRNAs assigned) and the number assigned an NTC gRNA (single-gRNA assignments only). **(D)** Bar plots showing the percentage of cells assigned no gRNA, a single gRNA, or multiple gRNAs. **(E)** Density plots of per-cell UMI distributions in cells assigned an NTC gRNA (grey) versus a targeting gRNA (blue). **(F)** Heatmap showing the fold-change in expression of all 240 TFs (rows) in cells assigned a gRNA for each of the 240 TFs (columns), relative to NTC control cells. The right-most column shows the fold-change in expression of all 240 TFs in NTC control cells relative to cells not assigned a gRNA. Values failing significance (adjusted p ≥ 0.05) were set to zero. **(G)** Violin plots showing the expression of each targeted TF in cells assigned the corresponding gRNA (blue, right), or the expression of all other murine TFs in NTC cells (grey, left), normalized to cell count. White boxplots indicate the median and interquartile range.

**Figure S5.**
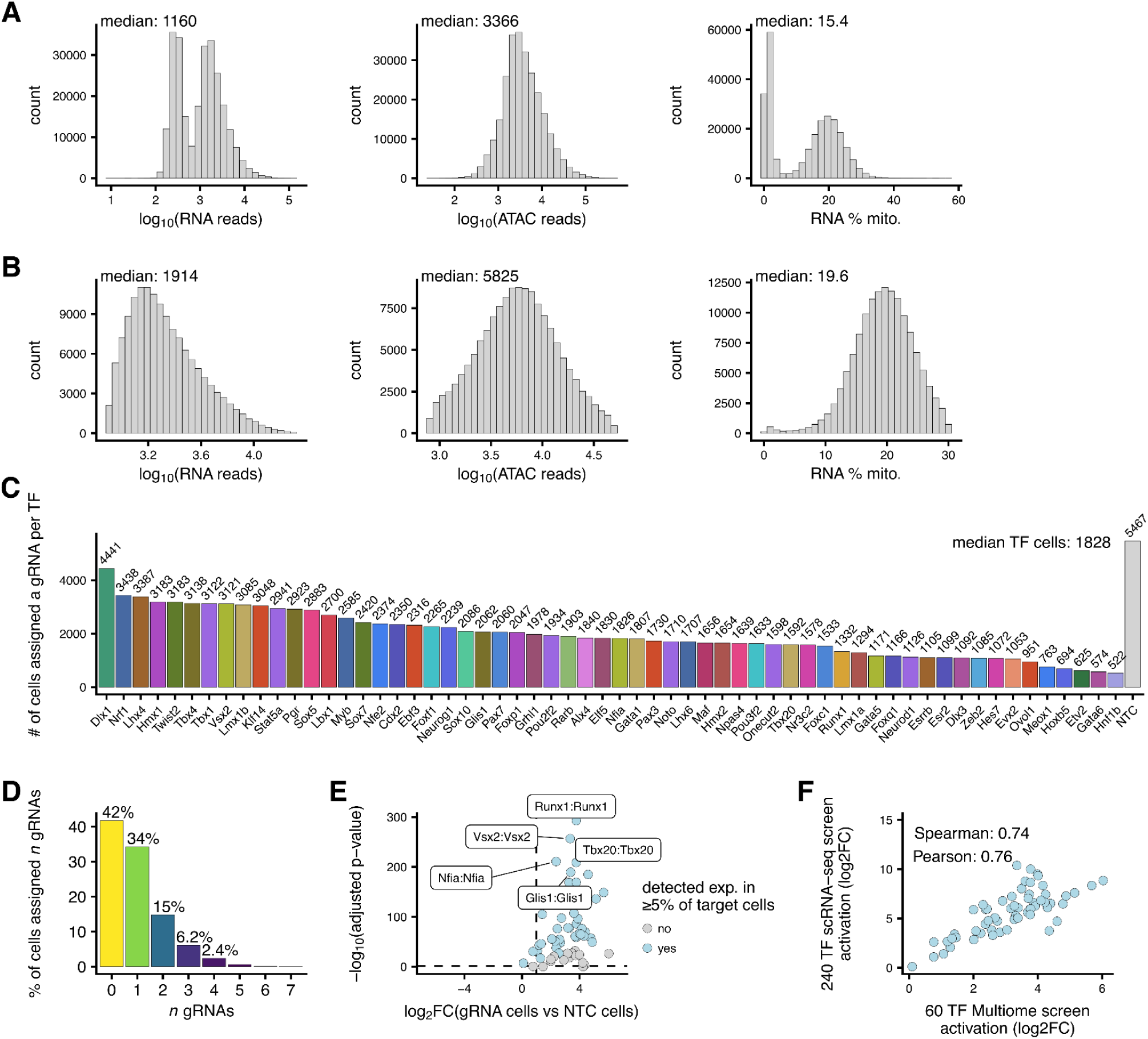
Cell filtering and gRNA capture for the 60-TF sc-multiome screen. **(A)** From left to right, distributions of per-cell RNA UMIs, unique ATAC fragments, and RNA mitochondrial read percentages, before filtering. **(B)** As in **A**, after filtering for cells with mitochondrial read percentage ≤30%, RNA and ATAC counts ≥800, ATAC counts <50,000 and RNA counts <20,000 (n = 139,374 cells). **(C)** Bar plots indicating the number of cells assigned a gRNA for each targeted TF (including cells with multiple gRNAs assigned) and the number assigned an NTC gRNA (single-gRNA assignments only). **(D)** Bar plots showing the percentage of cells assigned no gRNA, a single gRNA, or multiple gRNAs. **(E)** Fold change in expression of each target TF in cells bearing the corresponding gRNA versus NTC gRNA(+) cells. Fill indicates whether expression of the corresponding TF was detected in at least 5% of that TF’s gRNA(+) cells. **(F)** Correlation in target TF activation levels (log₂FC of gRNA(+) cells relative to NTC cells) between the 240-TF scRNA-seq screen (y-axis) and the 60-TF sc-multiome screen (x-axis).

**Figure S6.**
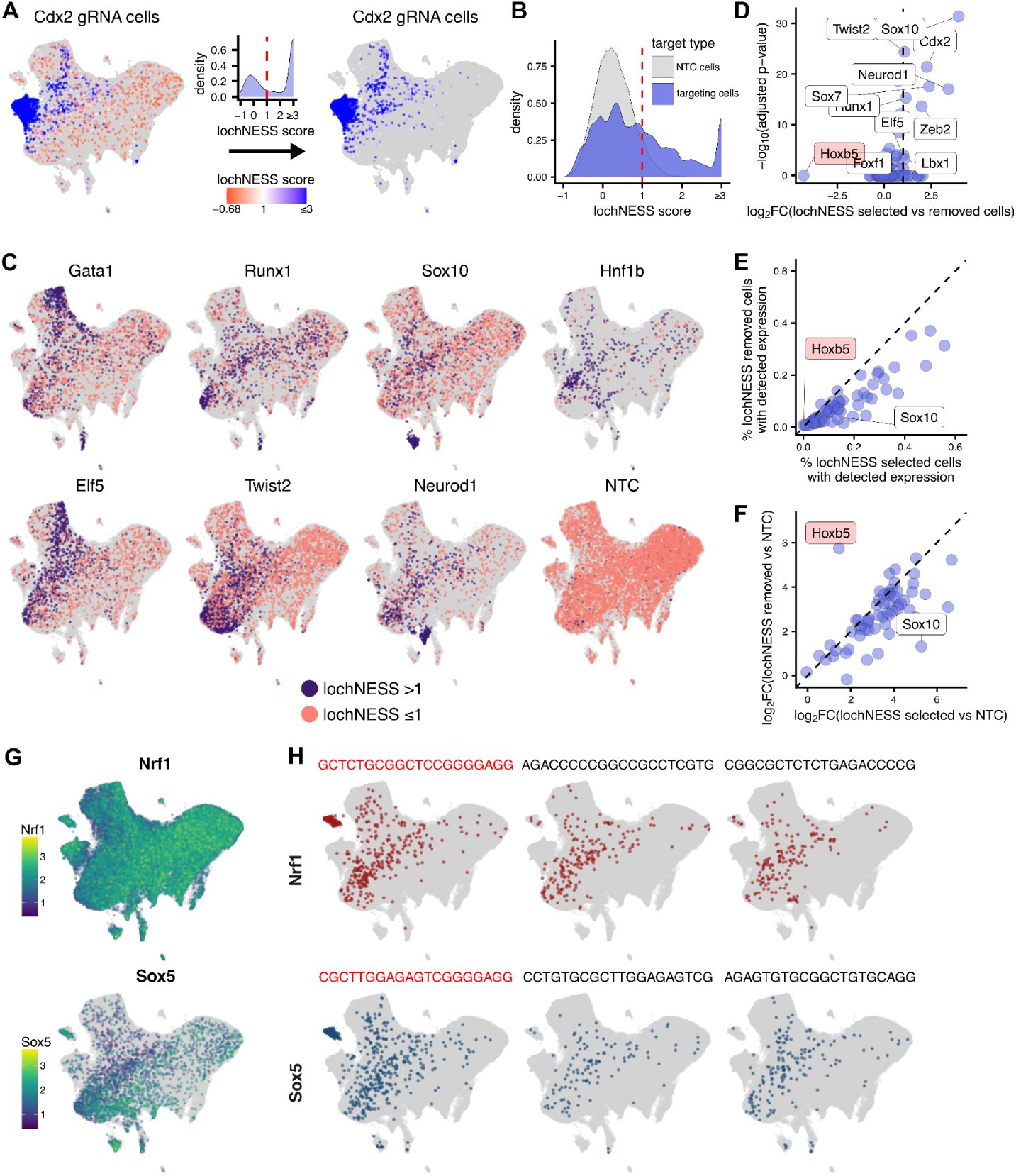
lochNESS filtering selects for more efficiently perturbed cells. **(A)** ATAC UMAP showing lochNESS scores for *Cdx2* gRNA(+) cells before (left) and after (right) applying a cutoff of 1. *Cdx2* gRNA(+) cells are colored by their lochNESS score; all other cells are light grey. **(B)** Density distributions of lochNESS scores for TF-targeting gRNA(+) cells (purple, pooled across TFs) and non-targeting control (NTC) gRNA(+) cells (grey). Scores were calculated separately and are pooled here for visualization, capping at 3; the dashed vertical line marks the cutoff of 1. **(C)** ATAC UMAPs for several targeting gRNAs and an NTC control. Purple, gRNA(+) cells retained by lochNESS (score > 1); red, gRNA(+) cells removed (score ≤ 1); light grey, all other cells. **(D)** Volcano plot of differential expression of the target TF between lochNESS-retained and lochNESS-removed gRNA(+) cells for that TF (x-axis, log₂ fold change; y-axis, −log_10_ adjusted p-value, Wilcoxon). Dashed vertical line, log_2_FC = 1. The top 10 most significant differences with log_2_FC > 1 are labeled. *Hoxb5* appears as a prominent outlier with a negative fold change, likely reflecting stochastic differences arising from the extremely low fraction of cells with detectable *Hoxb5* expression in both groups. **(E)** Scatter plot comparing, for each target TF (one point per TF), the percentage of lochNESS-retained (x-axis) and removed (y-axis) gRNA(+) cells with detected target TF expression. Dashed diagonal, x = y. **(F)** As in **E**, but comparing log₂ fold-change in target TF expression relative to NTC cells. Labels mark the TFs with the largest positive and most negative values of Δ(log_2_FC retained − log₂FC removed); *Hoxb5* is again an outlier, for the reason given in **D**. **(G)** Expression distributions of *Nrf1* (top) and *Sox5* (bottom) in the ATAC UMAP embedding. **(H)** ATAC UMAP distributions of cells assigned each of the three *Nrf1*-targeting gRNAs (top row) and each of the three *Sox5*-targeting gRNAs (bottom row). For each gRNA, colored points are cells that passed lochNESS filtering (score > 1); all others are grey. The gRNAs highlighted in red share sequence similarity and show a distinct distribution in UMAP space.

**Figure S7.**
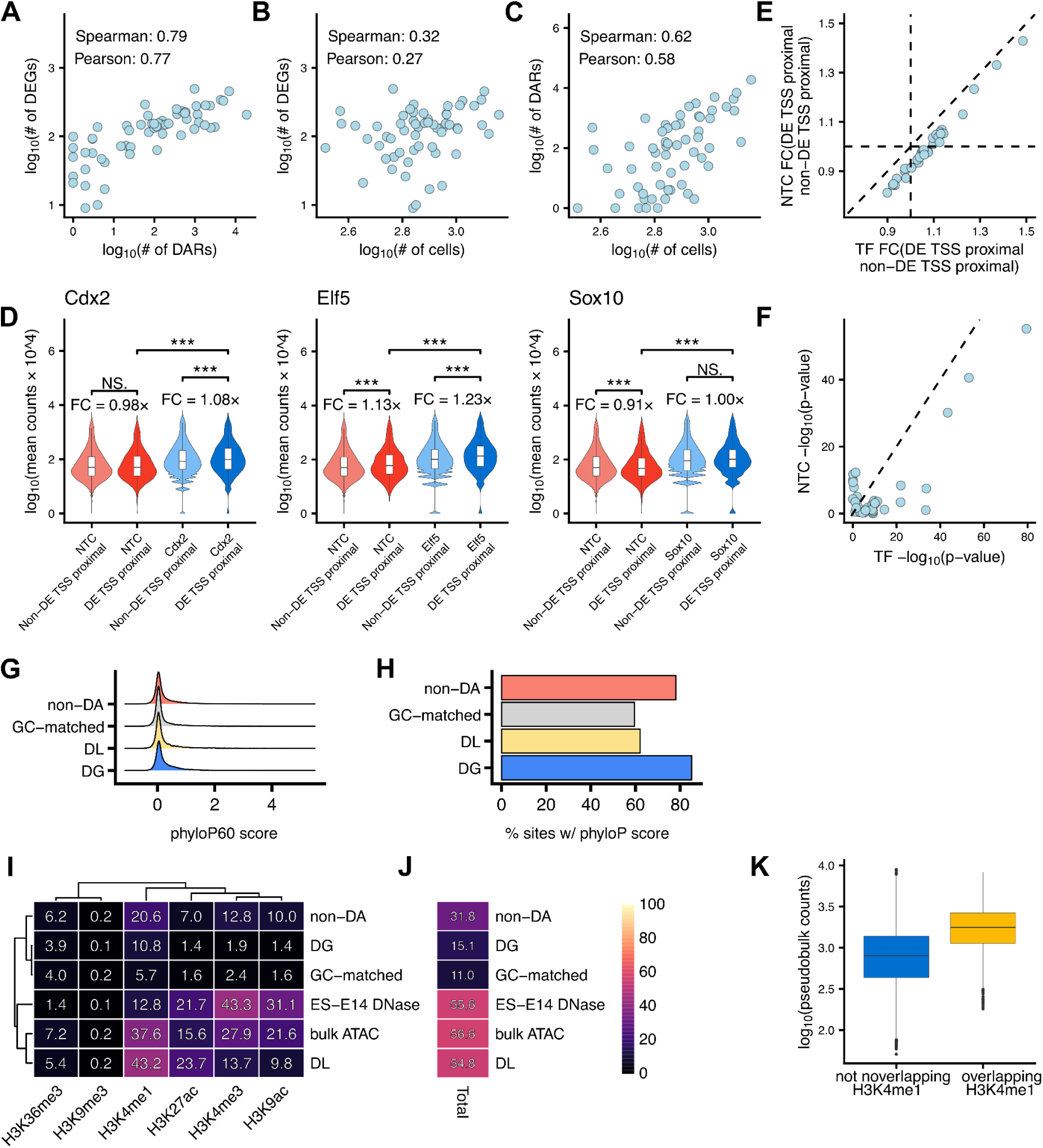
Characterizing differentially accessible regions. **(A)** Scatter plot showing the correlation between the number of DA sites and the number of DEGs across TFs. **(B)** Scatter plot showing the correlation between the number of cells assigned to each TF and the number of DEGs detected for that TF. **(C)** Scatter plot showing the correlation between the number of cells assigned to each TF and the number of DA sites called for that TF. **(D)** Increased accessibility at peaks near differentially expressed genes in TF-targeted cells. Violin plots show per-peak pseudobulk accessibility (mean ATAC counts per cell; plotted as log_10_(mean counts × 10^4^ + 1)) for peaks within ±100 kb of the TSS of genes differentially expressed by the indicated TF (“DE TSS proximal”) compared to peaks within ±100 kb of the TSS of all other genes with detected expression (“non-DE TSS proximal”). Values are shown for NTC cells (red) and lochNESS-selected TF cells (blue) for several representative TFs. Asterisks indicate Wilcoxon rank-sum test significance for the indicated comparisons. **(E)** Scatter plot showing, for each TF, the fold change (FC) in mean per-peak ATAC accessibility (mean counts per cell) for peaks within ±100 kb of TSSs of DE genes relative to peaks within ±100 kb of TSSs of non-DE genes. FC values are shown for NTC cells (y-axis) and for lochNESS-selected TF cells (x-axis). **(F)** As in **E**, showing the −log_10_ p-values (Wilcoxon rank-sum test) of the fold-change comparisons. **(G)** Peaks were categorized as differentially gained (DG), differentially lost (DL), or non-differentially accessible (non-DA), and an additional background set of random genomic regions was generated from mm10, matched to DG regions by size and GC composition. Density plots show the distributions of phyloP scores for each of these categories among peaks that had a phyloP score. **(H)** Bar plots showing the percentage of sites for each category defined in **G** that had a phyloP score. **(I)** The same peak categories as defined in **G**, here also including ENCODE peaks from ES-E14 DNase-seq data and bulk ATAC peaks called on our WT mESC CRISPRa cell line. The heatmap displays the fraction of peaks in each category (y-axis) that overlap ENCODE ChIP-seq peaks generated in serum-cultured E14 mESCs for each of the indicated histone modifications (x-axis). Differences in cell line and culture conditions (2i + LIF in the current dataset versus serum + LIF in the ENCODE datasets), and the shallower sequencing depth of the ChIP-seq datasets, could contribute to lower overlap percentages. The DNase dataset is also shallow (n = 62,535 peaks), contributing to its outgroup clustering. **(J)** Heatmap showing for each peak category the total fraction of peaks overlapping one or more ENCODE ChIP-seq peaks across the set of histone marks analyzed. **(K)** Boxplots showing log_10_-transformed total fragment counts for DG peaks, stratified by whether they overlap H3K4me1 ENCODE peaks in WT mESCs (10.9%, n = 3,011) or not (89.1%, n = 24,731), indicating that a subset of DG sites may already exhibit some accessibility in unperturbed mESCs.

**Figure S8.**
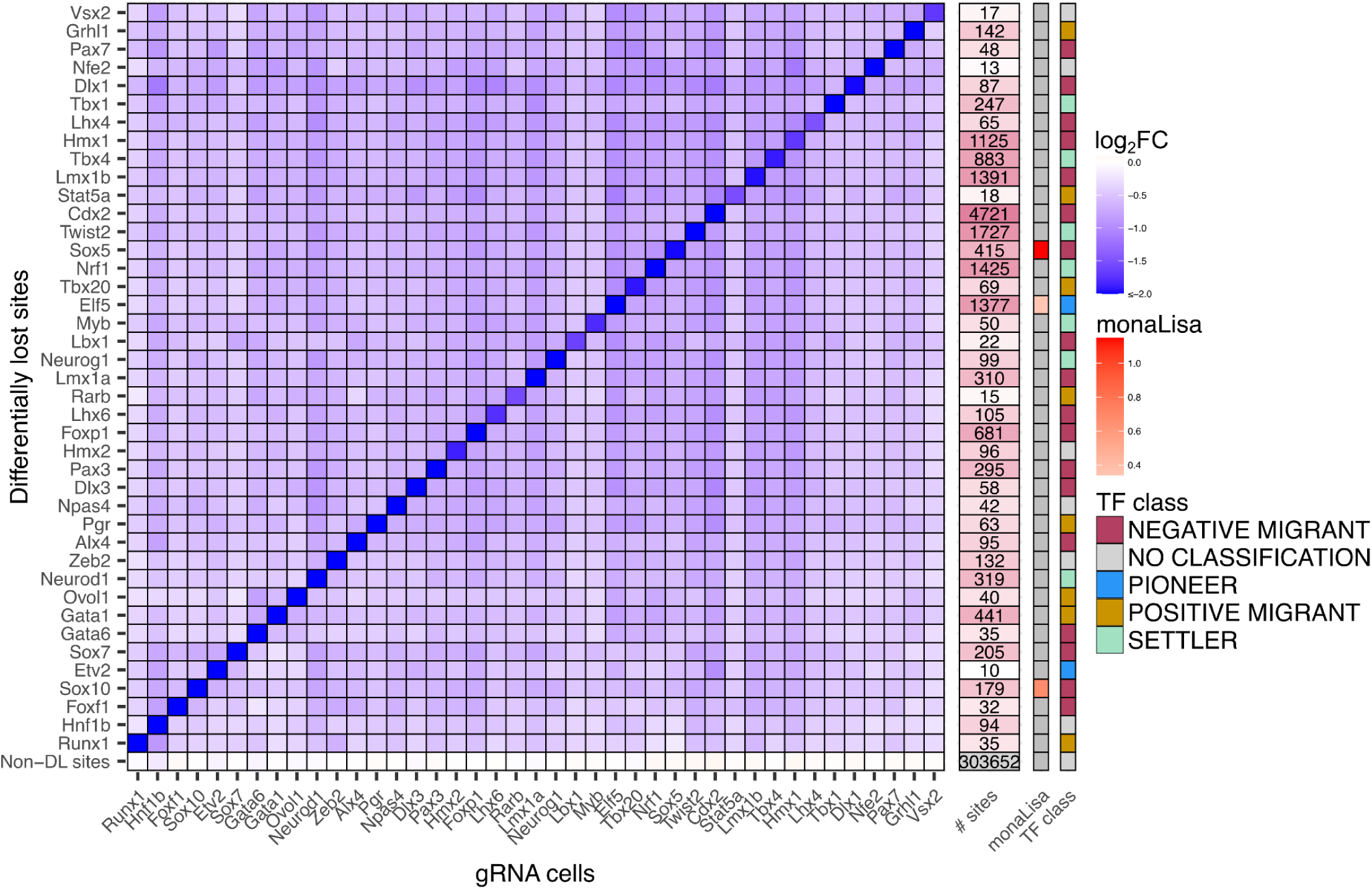
Overlap in chromatin accessibility changes at DL sites across targeted TFs. Heatmap showing the average log₂ fold-change in chromatin accessibility (gRNA vs NTC) at differentially lost (DL) sites defined for each TF (rows) across lochNESS-selected gRNA(+) cell populations (columns). DL sites were called for each TF using an empirical 5% FDR threshold; cross-TF entries reflect the average log_2_ fold-change at those sites, without reapplying significance criteria, capped at ±3 for visualization. TFs (rows) were hierarchically clustered (Ward.D2) on their DL-site accessibility profiles. Only TFs with at least 10 DL sites are shown. Non-DL sites (top row) correspond to the average log_2_ fold-change across all peaks not called DL for any TF. Adjacent vertical tile panels indicate, from left to right, the number of DL sites contributing to each TF’s row; the enrichment of the matching TF motif family within DL sites, calculated using monaLisa with GC-matched genomic background sequences (most enriched motif from the matching TF family with −log_10_ adjusted p > 2), and the chromatin-opening classification assigned to each TF by Ehsani *et al.*^45,46^

**Figure S9.**
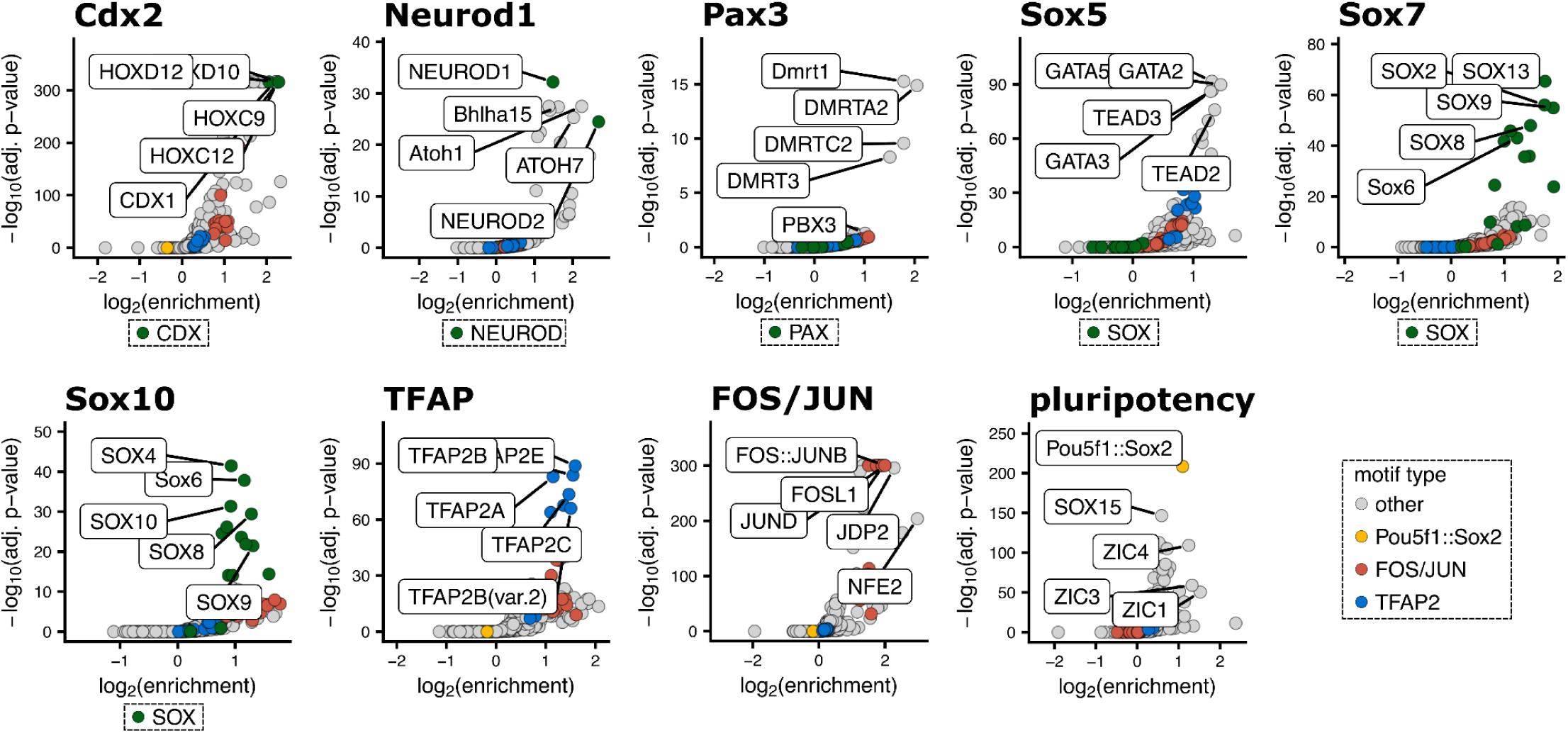
Motif enrichment in DA peak types. Motif enrichment within each group of DA sites: (i) six TF-specific groups (*Cdx2*, *Neurod1*, *Pax3, Sox5, Sox7*, and *Sox10*), comprising sites called DG exclusively for the indicated TF (for *Sox5*, including sites DG for *Sox5* and/or *Nrf1* but no other TF); (ii) TFAP, comprising sites DG for one or more trophoblast-associated TFs (*Elf5*, *Grhl1*, *Gata1*, *Gata5*, *Gata6*, *Ovol1*) and no other TFs; (iii) FOS/JUN, comprising remaining DG sites not assigned to the above categories; and (iv) pluripotency, comprising sites called DL for any TF and DG for none. For each group, motif enrichment was performed with monaLisa using GC-matched genomic background sequences. Volcano plots show log_2_(enrichment) on the x-axis and −log_10_(adjusted p-value) on the y-axis. In the color legend, motifs are classified as “FOS/JUN” if the motif ID contains “Fos” or “Jun”, as “TFAP2” if it contains “TFAP2”, as “Pou5f1::Sox2” for that specific motif, and as “other” for all remaining motifs. For TF-specific groups, motifs whose IDs match the target TF family name (*e.g.* NEUROD for *Neurod1*, SOX for *Sox7*) were additionally assigned to a TF-family-specific category and colored separately (green). The top 10 motifs by combined enrichment and significance score (log₂(enrichment) × −log_10_(adjusted p-value)) are labeled.

**Figure S10.**
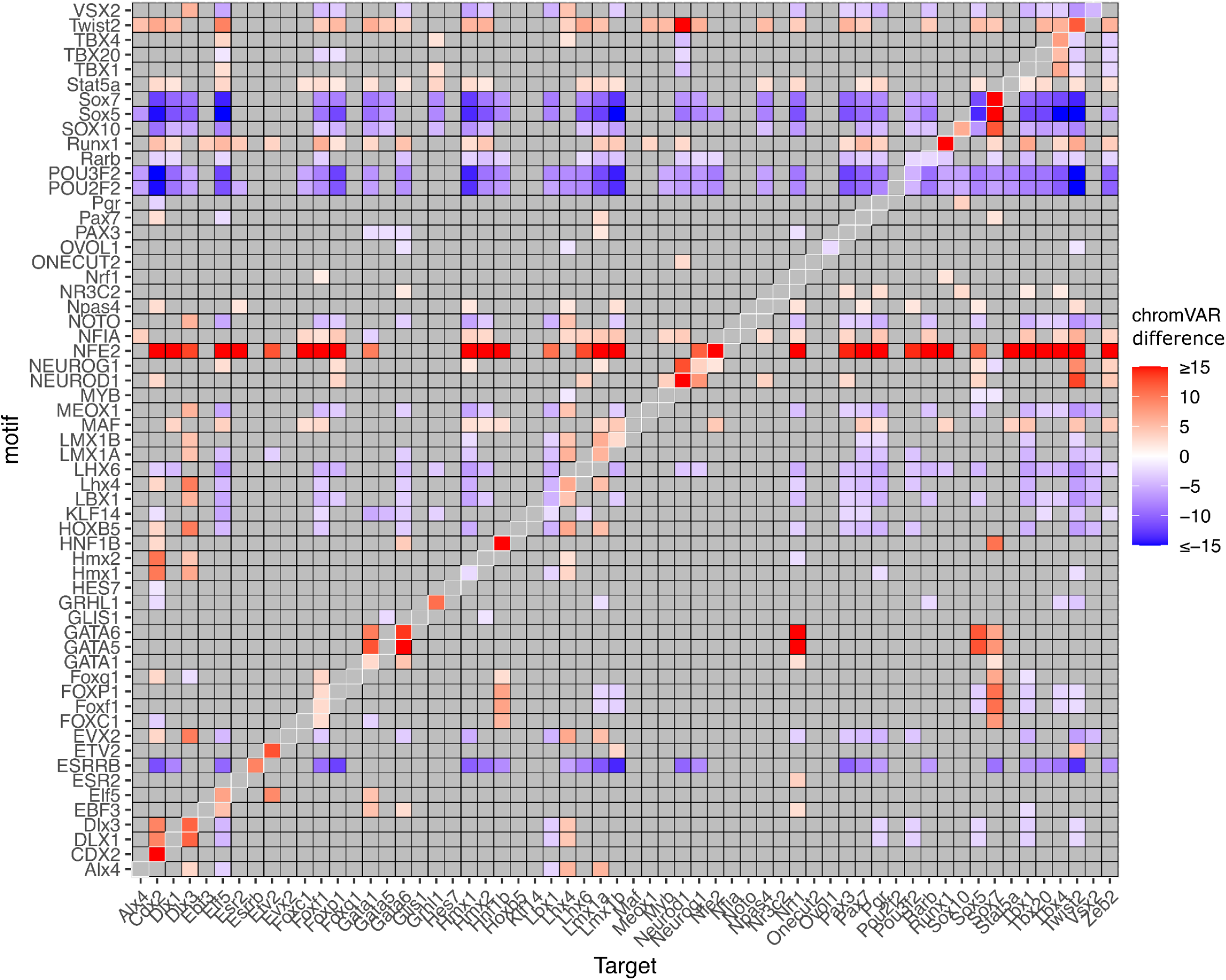
Target TF motif accessibility analysis with chromVAR. Heatmap showing the difference in chromVAR-derived motif accessibility in target TF gRNA(+) cells (columns) relative to NTC gRNA(+) cells, for each TF’s exact motif match (rows), not showing related motif family members as in **Fig. 3D**. Tiles are colored by the chromVAR differences when adjusted p < 0.05, with nonsignificant values (adjusted p ≥ 0.05) shown in grey. White boxes along the diagonal highlight each TF’s own motif, of which 21 show significantly increased accessibility (adjusted p < 0.05; chromVAR difference > 0).

**Figure S11.**
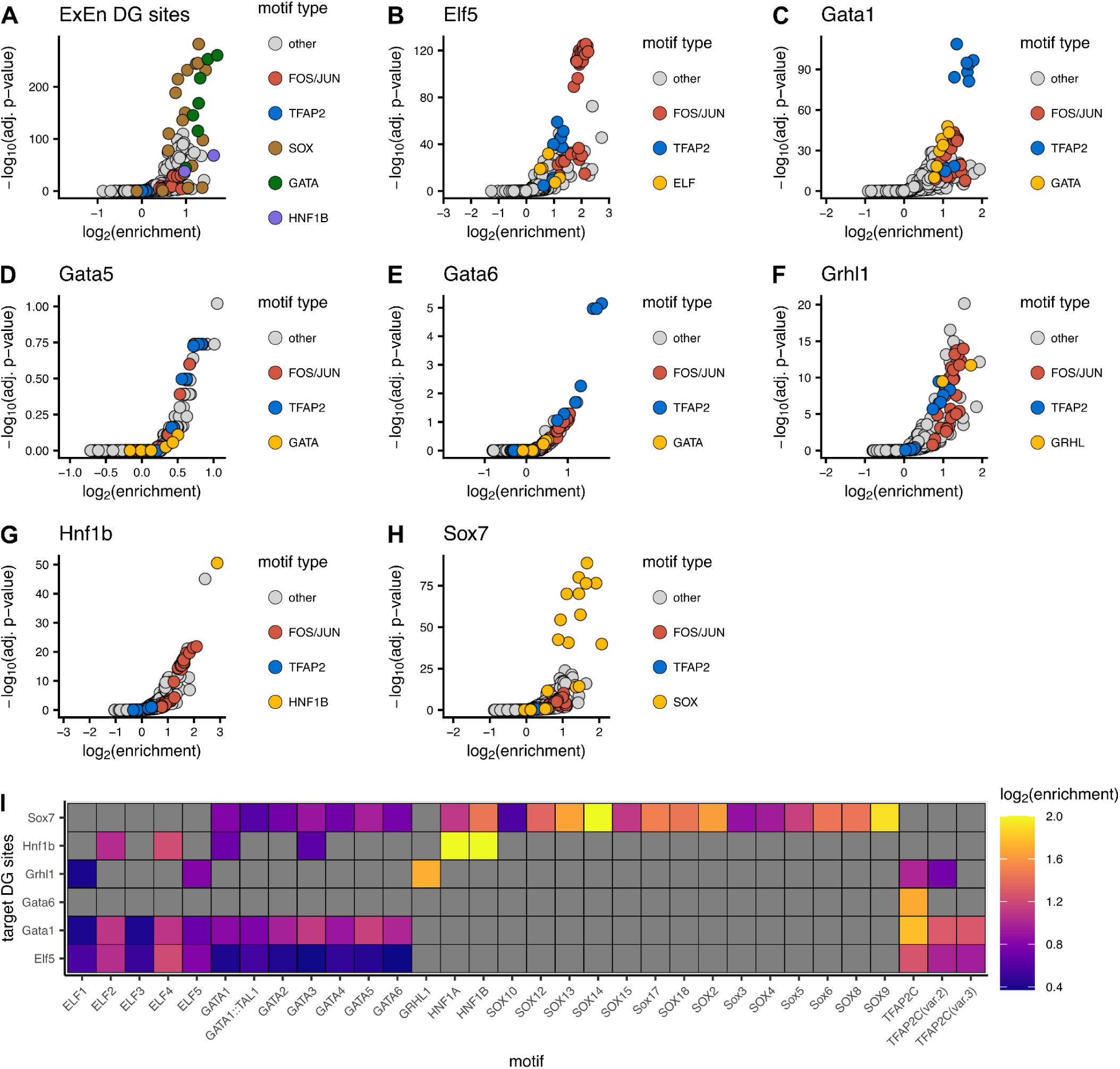
Motif enrichment of target TF motifs and the TFAP2C motif in TF DG sites. **(A)** Motif enrichment in DG sites called on all cells within the ExEn-annotated ATAC cluster. **(B-H)** Motif enrichment in the DG sites of each indicated TF. In **A-H**, enrichment was performed with monaLisa using GC-matched genomic background sequences, and volcano plots show log_2_(enrichment) on the x-axis and −log_10_(adjusted p-value) on the y-axis. Motifs are colored as “FOS/JUN” if the motif ID contains “Fos” or “Jun”, “TFAP2” if it contains “TFAP2”, as the TF-specific family (*e.g.* GATA, SOX) for motifs matching the indicated TF, and “other” for all remaining motifs. **(I)** Heatmap showing motif enrichment, computed as in **A-H**, for the ELF, GATA, GRHL, HNF1, SOX and TFAP2 motif families in the DG sites of *Elf5*, *Gata1*, *Gata6*, *Grhl1*, *Hnf1b* and *Sox7* gRNA(+) cells.

**Figure S12.**
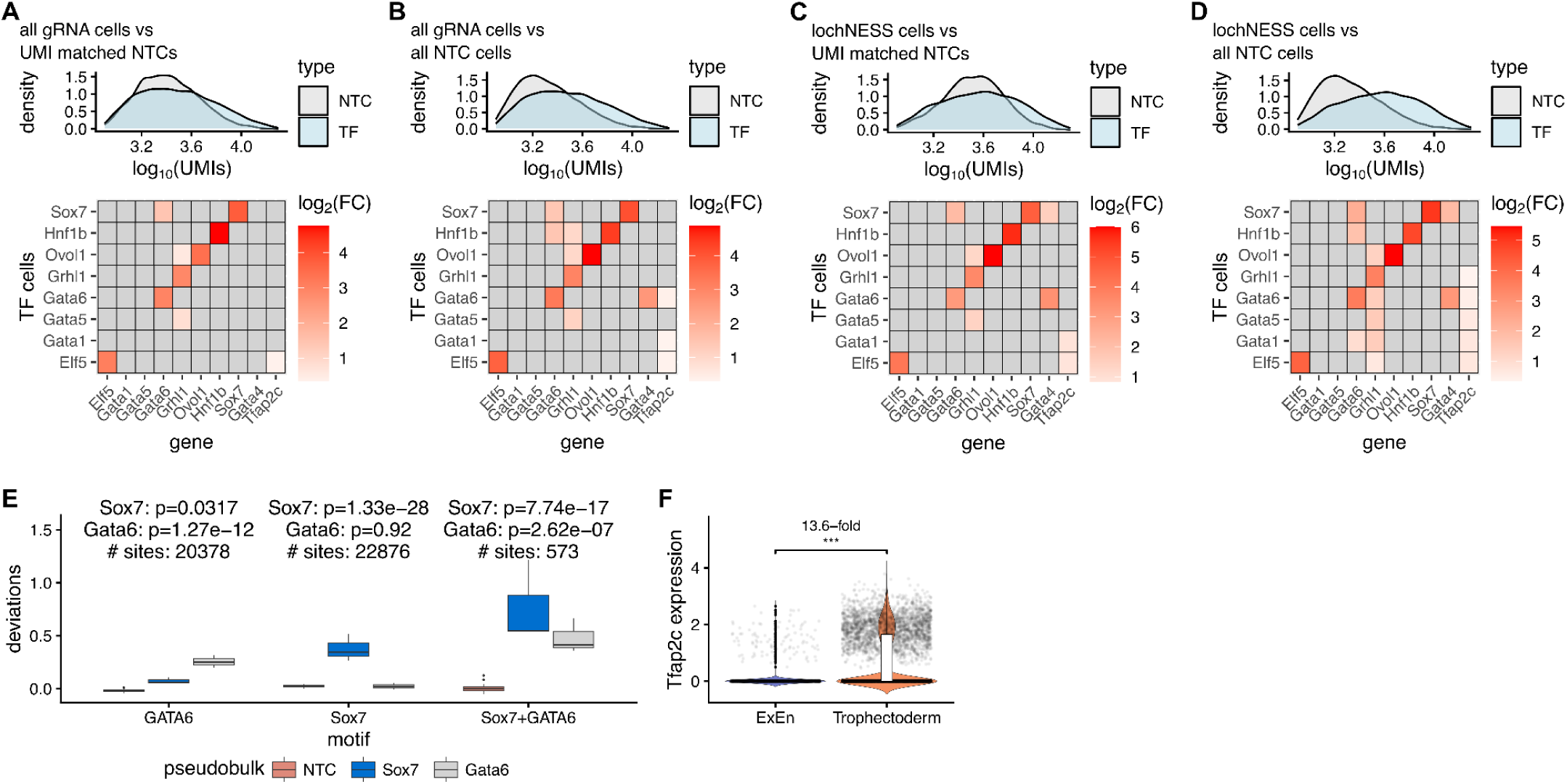
Differential expression comparing TF gRNA(+) cells to NTC cells under different cell selection strategies. Heatmaps (lower subpanels) show differential expression of the indicated genes (columns) in TF gRNA(+) cells (rows) relative to NTC cells. TF cells were either all gRNA(+) cells or lochNESS-selected gRNA(+) cells, and NTC cells were either all NTC cells or NTC cells subsampled to match the per-cell UMI distribution of TF cells (as indicated in panel titles). Differential expression was tested by Wilcoxon rank-sum test. Values are grey if they failed BH-adjusted p < 0.01, showed <1.2-fold increase, or were detected in <5% of TF cells. Density plots (upper subpanels) show the log_10_-normalized per-cell UMI distribution of the TF cells used in each comparison (blue) and of the corresponding NTC cells (light grey). In all comparisons, *Gata1* and *Gata5* showed significant activation (≥4-fold increase; BH-adjusted p < 0.01) but appear grey along the diagonal because the target transcript was detected in <5% of TF cells. **(A)** All TF gRNA(+) cells vs UMI-matched NTC cells; an increase in *Tfap2c* expression passing thresholds was observed only for *Elf5* (1.27-fold). **(B)** All TF gRNA(+) cells vs all NTC cells; the largest such *Tfap2c* increase was for *Gata6* (1.31-fold). **(C)** lochNESS-selected TF gRNA(+) cells vs UMI-matched NTC cells; the largest such *Tfap2c* increase was for *Gata1* (1.83-fold). **(D)** lochNESS-selected TF gRNA(+) cells vs all NTC cells; the largest such *Tfap2c* increase was for *Elf5* (1.62-fold). **(E)** Boxplots of chromVAR deviations for GATA6 motif, SOX7 motif, and SOX7+GATA6 co-motif instances (motifs within 5-100 bp of one another) in pseudobulked *Gata6*, *Sox7* and NTC cells. Adjusted p-values for *Sox7* vs NTC and *Gata6* vs NTC are indicated above each pair of boxplots, along with the number of motif instances used. **(F)** Violin plots of *Tfap2c* expression in cells from the ExEn-like cluster (left) and the trophectoderm-like cluster (right), with fold change and significance for the comparison between clusters.

**Figure S13.**
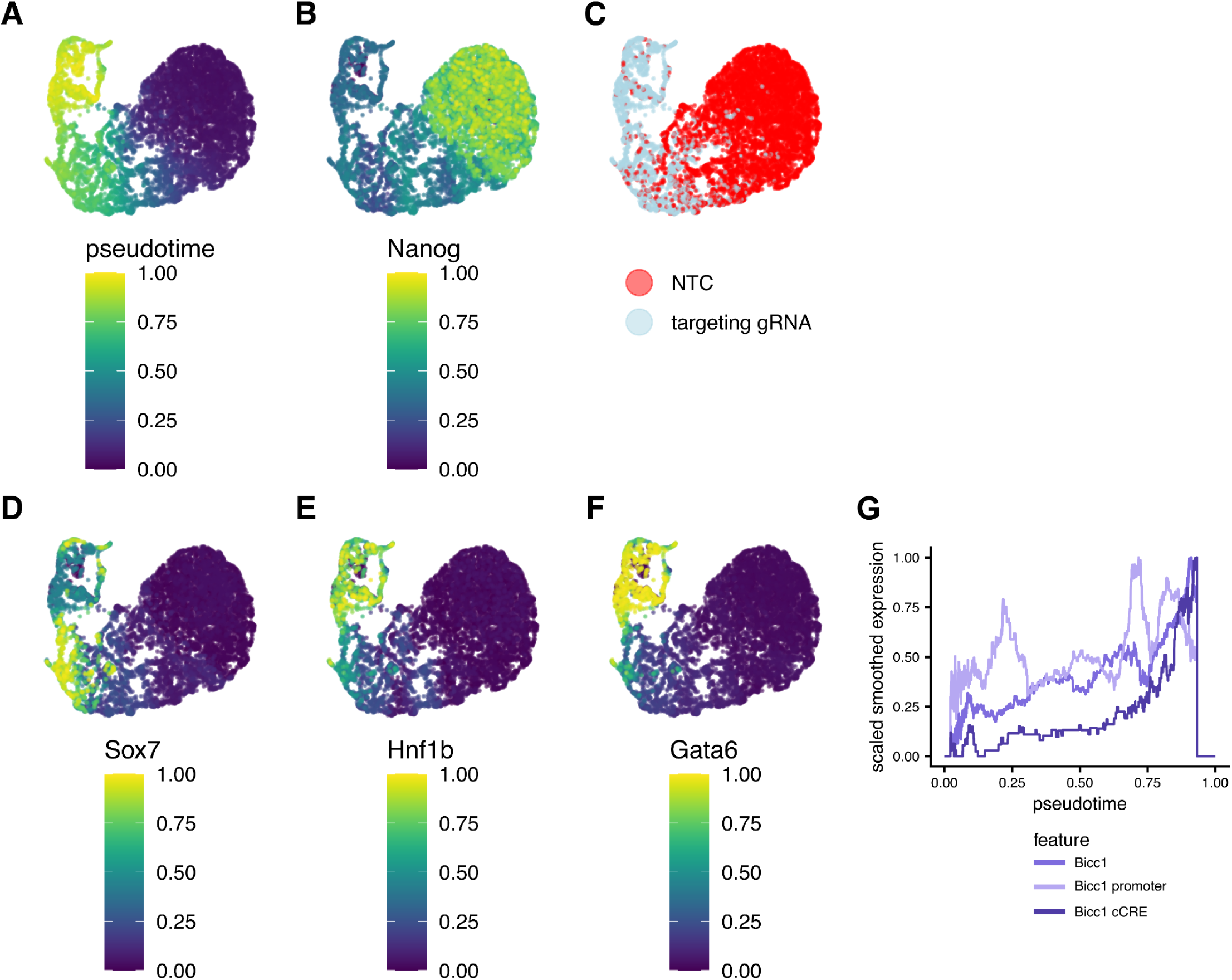
Pseudotime ordering of cells within the ExEn-like state. **(A)** UMAP visualization of the diffusion pseudotime embedding, constructed using all cells from the ExEn-like cluster together with NTC gRNA(+) cells. Pseudotime values inferred from the diffusion map are overlaid onto the embedding. **(B)** Same embedding, showing smoothed expression of *Nanog*. (**C)** Same embedding, showing the distribution of NTC gRNA(+) cells versus all targeting gRNA(+) cells. **(D-F)** Smoothed expression profiles across pseudotime for *Sox7* **(D)**, *Hnf1b* **(E)** and *Gata6* **(F)**. **(G)** Smoothed expression and accessibility profiles across pseudotime for *Bicc1*, for a peak overlapping the *Bicc1* promoter, and for a peak intronic to *Bicc1* (“*Bicc1* cCRE”). Throughout, expression and accessibility values were smoothed by rolling average across neighboring cells ordered by pseudotime and scaled from 0 to 1 independently for each feature.

**Figure S14.**
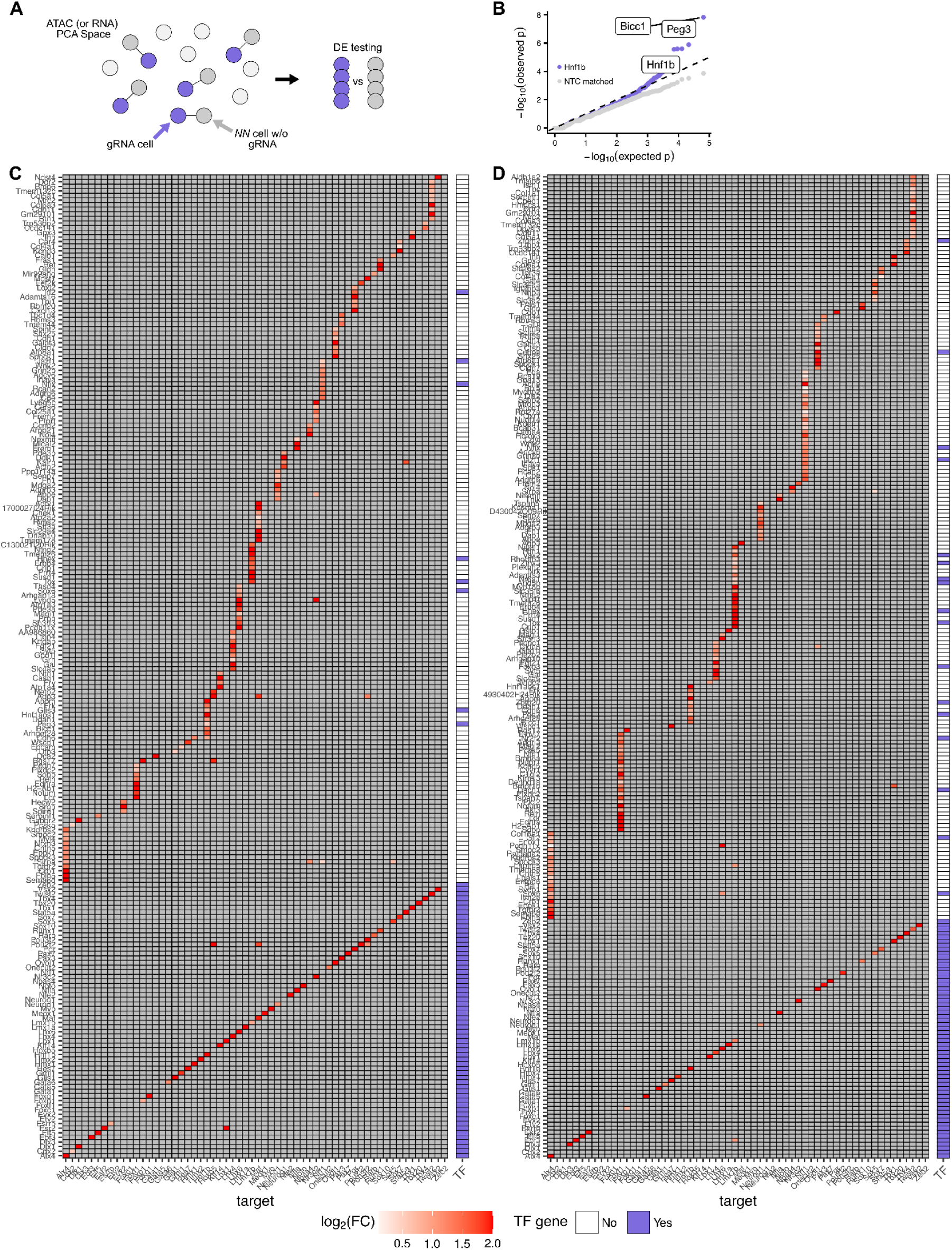
Nearest-neighbor (NN) differential expression across all TF perturbations. **(A)** Schematic representation of NN approach. For each gRNA cell the NN cell in PC space is identified and used as out group for differential expression (or accessibility) testing. (**B**) Quantile-quantile plot showing distribution of expected vs observed p-values from running the NN DE testing on *Hnf1b* gRNA(+) cells (purple dot) or creating a null distribution by performing the same analysis using a matched number of NTC cells (grey dots). **(C)** Heatmap summarizing NN differential expression analysis for all TF perturbations using all gRNA-assigned cells. For each TF-targeting gRNA, cells were matched one-to-one with the nearest cell in ATAC PCA space that was assigned either a different TF-targeting gRNA or a non-targeting control (NTC) gRNA and had not already been selected as a control. This matching procedure minimizes differences arising from pre-existing chromatin state heterogeneity while excluding cells assigned the same perturbation. Differential expression was then performed between each TF perturbation and its matched controls. Statistical significance was determined using an empirical false discovery rate (FDR) framework in which the NN differential expression analysis was repeated five times using matched NTC cells to generate an empirical null distribution of p-values. Empirical p-values were calculated as the fraction of null p-values at least as significant as the observed p-value, and an empirical FDR threshold of 10% was determined for each TF using the Benjamini–Hochberg procedure applied to these empirical p-values. Genes with average log_2_ fold change >0.1 passing the corresponding empirical FDR threshold are shown in color, whereas non-significant comparisons are shown in gray. Rows include all targeted TFs together with genes that were significantly upregulated for at least one perturbation. Colors indicate average log_2_ fold change relative to matched controls, with truncation of values greater than 2. The adjacent binary annotation indicates whether each gene is a curated murine transcription factor. **(D)** Same analysis as in **C**, except NN differential expression was performed using only the subset of cells assigned to each perturbation and passing lochNESS filtering.

**Figure S15.**
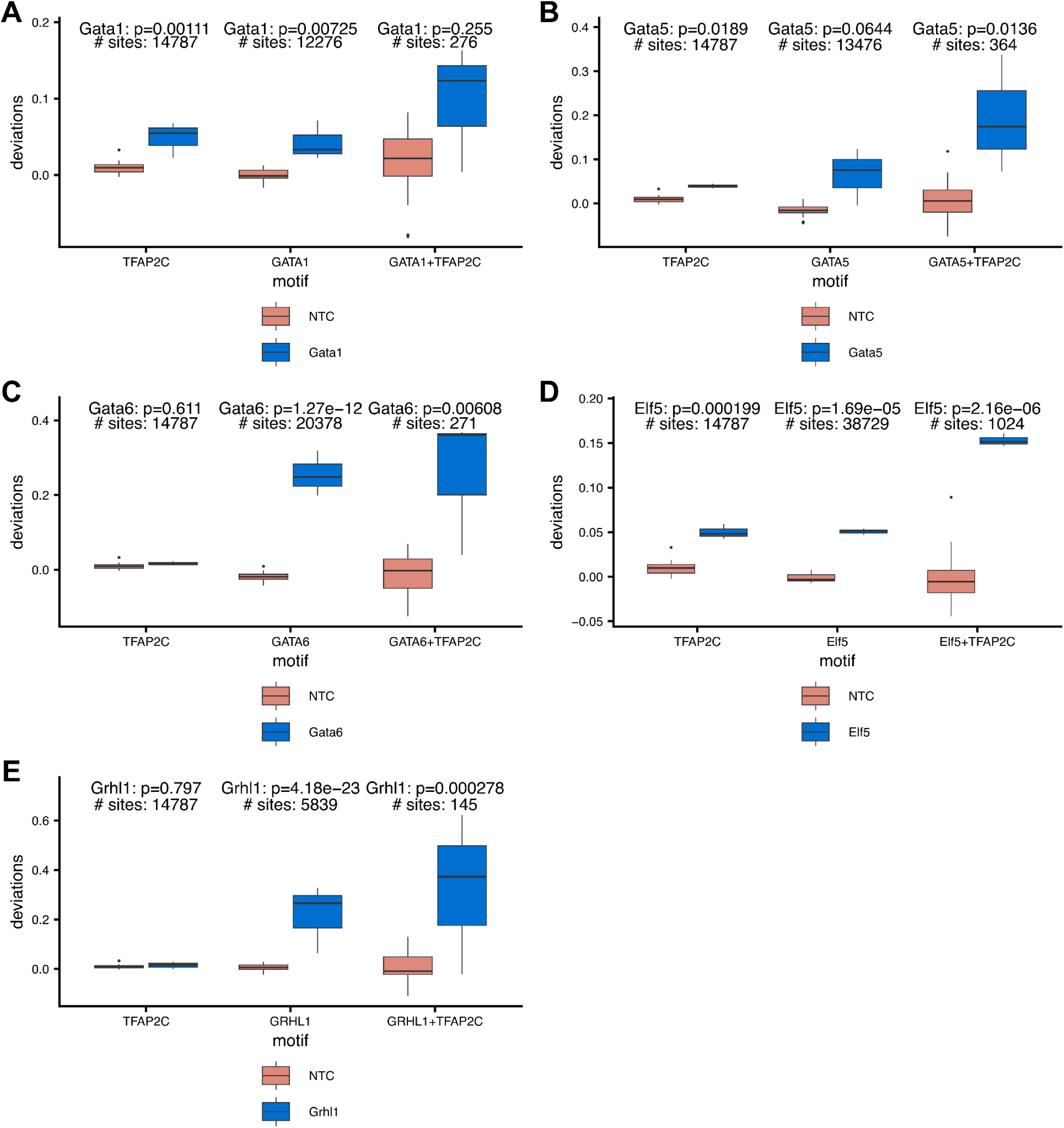
Increase in co-motif accessibility for different TF targets. **(A)** Boxplots showing chromVAR deviations for the TFAP2C motif, GATA1 motif, and GATA1–TFAP2C co-motif instances (in which the motifs are between 5-100bp of one another) for *Gata1* and NTC pseudobulked cells. The significance between *Gata1* and NTC cells is indicated above each pair of boxplots, as well as the number of sites / motif instances used for the analysis. **(B-E)** Same as **A**, shown for the indicated TF and its corresponding motif and TFAP2C co-motif classes.

**Figure S16.**
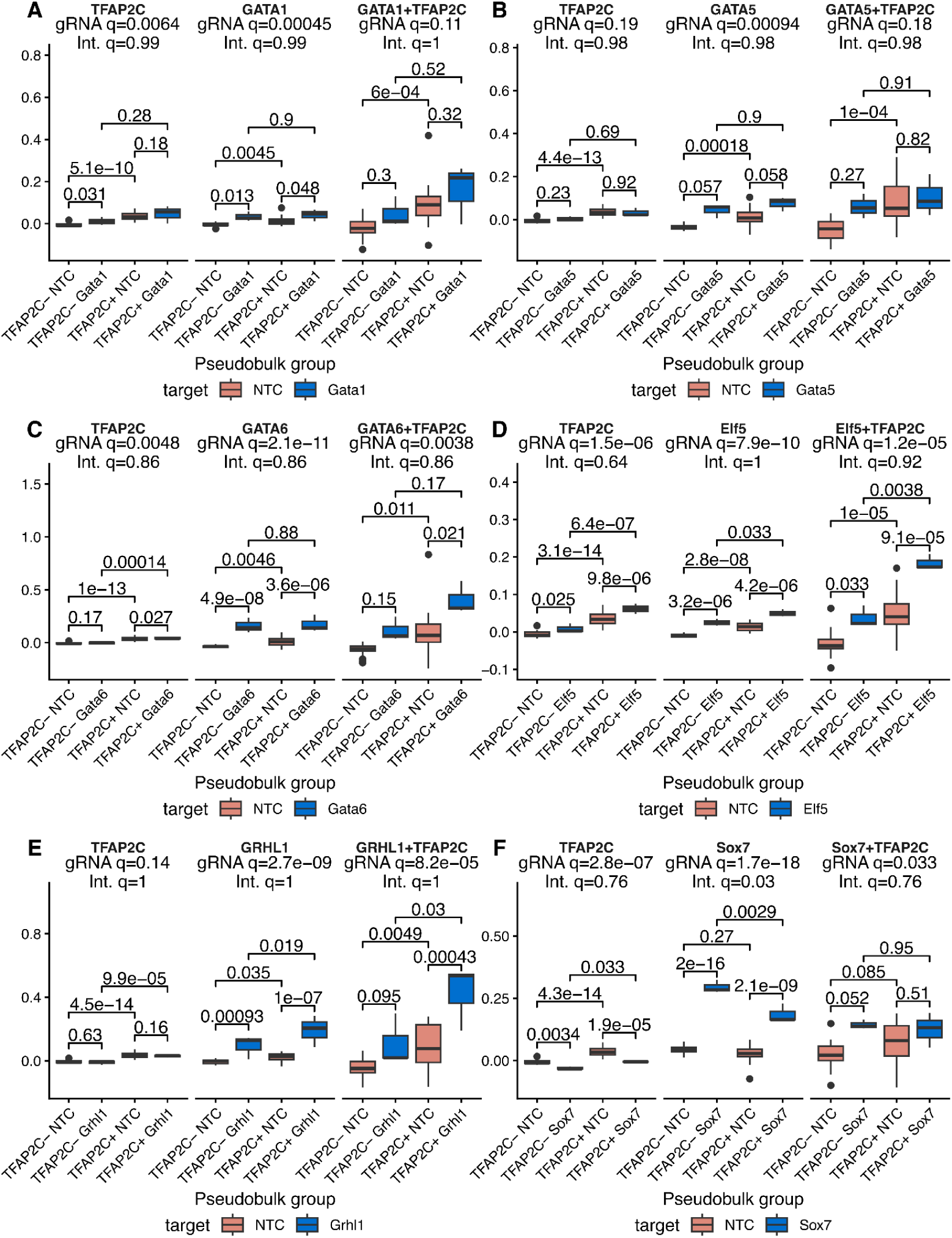
Co-motif enrichment for different TF targets after stratifying by *Tfap2c* expression (including cells with multiple gRNAs assigned). **(A**) Boxplots showing chromVAR deviation scores at TFAP2C motif sites, GATA1 motif sites, and GATA1–TFAP2C co-motif sites (motifs located within 5–100 bp of each other) in pseudobulked *Gata1* gRNA and NTC cells, stratified by detection of *Tfap2c* expression (TFAP2C+ and TFAP2C− cells). Adjusted p-values for between-group comparisons are shown. Above each motif class, the significance of the marginal (“global”) CRISPRa effect averaged across TFAP2C strata is indicated (“gRNA q =”), along with the significance of the CRISPRa × TFAP2C interaction from a limma model (“Int. q =”). **(B-F)** Same as **A**, shown for the indicated TF and its corresponding motif and TFAP2C co-motif classes.

**Figure S17.**
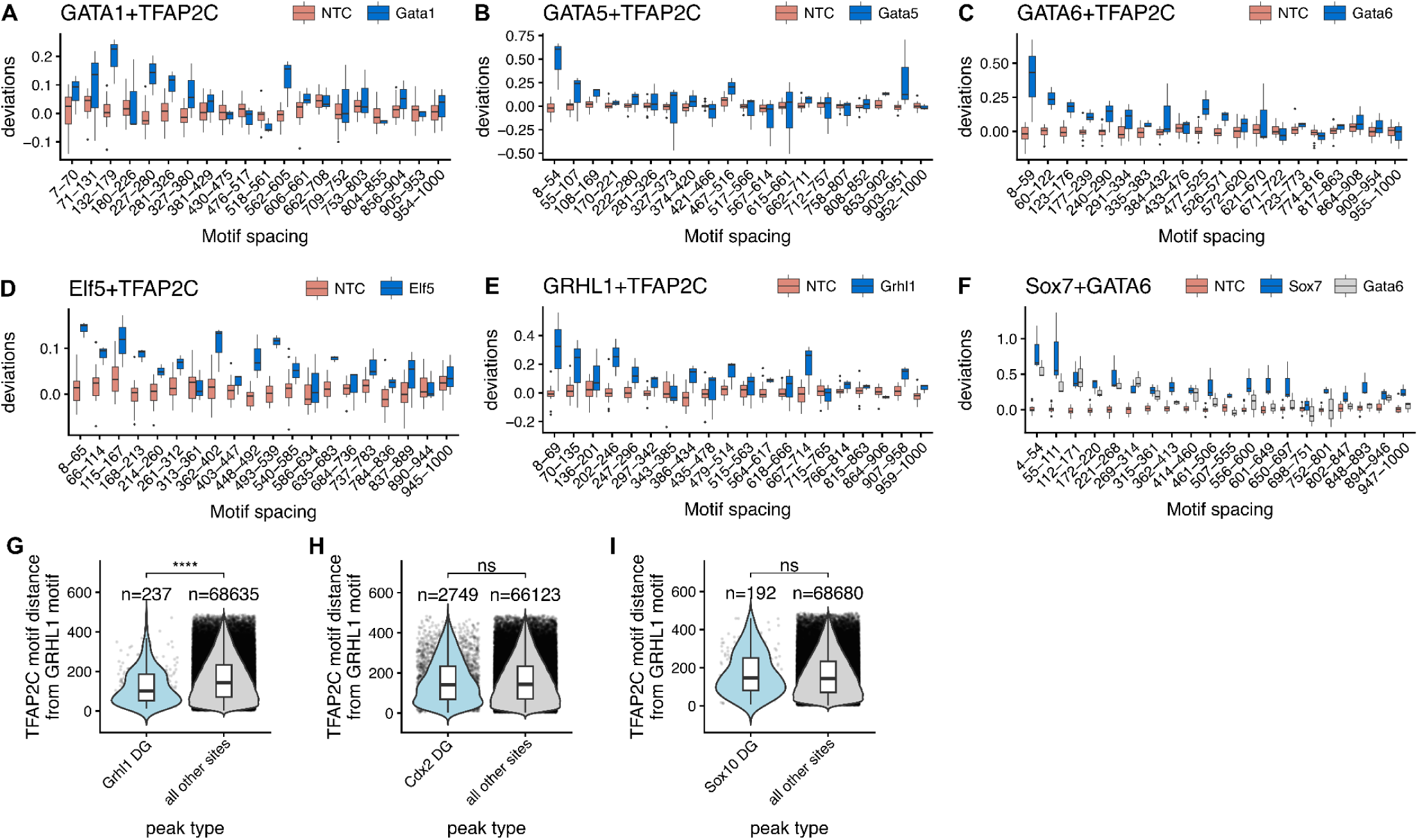
Co-motif enrichment for different TF targets across varying motif distance bins. (A-F) chromVAR deviation scores for genomic regions containing pairs of motifs, stratified by the distance between motif instances. For each transcription factor (TF) perturbation, accessibility deviations are shown for regions containing both motifs binned by inter-motif distance (x-axis; bins defined by quantiles of genomic distances between motif pairs, with bin ranges indicated). Boxplots display the distribution of deviation scores across pseudobulked cells for each condition, including TF-targeting gRNA(+) cells and non-targeting controls (NTC). Panels correspond to the following motif pairs: (**A**) The GATA1–TFAP2C co-motif in *Gata1* gRNA(+) cells, (**B**) The GATA5–TFAP2C co-motif in *Gata5* gRNA(+) cells, (**C**) The GATA6–TFAP2C co-motif in *Gata6* gRNA(+) cells, (**D**) The ELF5–TFAP2C co-motif in *Elf5* gRNA(+) cells, (**E**) The GRHL1–TFAP2C co-motif in *Grhl1* gRNA(+) cells, and (**F**) The Sox7–GATA6 co-motif in *Sox7* (blue) and *Gata6* (grey) gRNA(+) cells. **(G)** Motif spacing analysis between the TFAP2C motif (JASPAR MA0524.3) and the GRHL1 motif (JASPAR MA0647.2). All ATAC peaks were scanned using FIMO, retaining only the highest-scoring motif instance per peak for each motif (minimum FIMO p-value per peak). Peaks containing both motifs were identified, and the distance between motif midpoints was calculated. Violin plots show the distributions of GRHL1–TFAP2C motif distances in *Grhl1* differentially gained (DG) peaks (blue violins) versus all other peaks called on the dataset that contain both motifs (grey violins). Numbers above violins indicate the number of peaks containing both motifs per group. Statistical significance was assessed using a two-sided Wilcoxon rank-sum test. **(H)** Same as in **G**, here only looking at GRHL1–TFAP2C motif distances in *Cdx2* differentially gained sites, to control for differences being driven by how DG sites are called. **(I)** Same as in **G**, here only looking at GRHL1–TFAP2C motif distances in *Sox10* differentially gained sites, to control for differences being driven by how DG sites are called.

